# Deciphering complex interactions between LTR retrotransposons and three *Papaver* species using LTR_Stream

**DOI:** 10.1101/2025.03.30.646164

**Authors:** Tun Xu, Stephen J. Bush, Yizhuo Che, Huanhuan Zhao, Tingjie Wang, Peng Jia, Songbo Wang, Peisen Sun, Pengyu Zhang, Shenghan Gao, Yu Xu, Chengyao Wang, Ningxin Dang, Yong E. Zhang, Xiaofei Yang, Kai Ye

## Abstract

Long terminal repeat retrotransposons (LTR-RTs), a major type of class I transposable elements (TEs), are the most abundant repeat element in plants. The study of the interactions between LTR-RTs and the host genome relies on high-resolution characterization of LTR-RTs. However, for non-model species, this remains a challenge. To address this, we developed LTR_Stream for sub-lineage clustering of LTR-RTs in specific or closely related species, providing higher precision than current database-based lineage-level clustering. Using LTR_Stream, we analyzed Retand LTR-RTs in three *Papaver* species. Our findings show that high-resolution clustering reveals complex interactions between LTR-RTs and the host genome. For instance, we found autonomous Retand elements could spread among the ancestors of different subgenomes, like retroviruses pandemics, enriching genetic diversity. Additionally, we identified that specific truncated fragments containing transcription factors motifs such as TCP and bZIP may contribute to generation of novel TAD-like boundaries. Notably, our pre- and post-allopolyploidization comparisons show that these effects diminished after allopolyploidization, suggesting that allopolyploidization may be one of the mechanisms by which *Papaver* species cope with LTR-RTs. We demonstrated the potential application of LTR_Stream and provided a reference case for studying the interactions between LTR-RTs and the host genome in non-model plant species.

## INTRODUCTION

Transposable elements (TEs), also known as transposons, are mobile DNA elements that can cut or copy and insert themselves into different locations within a genome. In humans and other model organisms, extensive research has led to well-established TE classification systems [1]. For instance, human Alu elements can be categorized into 213 sub-subfamilies [2]. This refined classification forms the foundation for studying their complex evolutionary history [2] and interactions with host genomes [3–6]. In plants, long terminal repeat retrotransposons (LTR-RTs) are the most common type of TE [7], contributing significantly to the diversity and size variations observed between plant genomes [8–10]. However, unlike in humans and other model organisms, the fine-scale classification of transposons in non-model plants is still challenging, hindering further investigation into the impacts of specific TE types on host genomes. Especially in recent times, with advancements in sequencing and assembly technologies [11], researchers are generating an increasing number of complete plant genomes [8,12–16], leading to surging demanding for precise classification of LTR-RTs.

For example, our previously published genomes of three *Papaver* species, *P. rhoeas, P. somniferum*, and *P. setigerum* [13], recently underwent varying degrees of allopolyploidization, and contain one, two, and four subgenomes, respectively. These subgenomes have undergone complex processes of fission and fusion. However, as LTR-RTs constitute the largest proportion of sequences in these genomes [13], their intricate interactions, with host genomes under such a complex evolution scenario, remain unclear. As non-model organisms, current methods for classifying transposons, such as DeepTE [17] and TEsorter [18], typically categorize LTR-RTs into broad groups based on their family or lineage (lineage level), which lack the resolution to uncover subgenome-specific LTR-RTs. To address this, we developed LTR_Stream, a tool for fine-scale clustering of LTR-RTs in specific or closely related species. LTR_Stream represents LTR-RTs by identifying conserved sequence fragments (modules) within homology-based networks and clusters them using a graph-based learning approach. While constructing phylogenetic trees based on conserved protein domains within LTR-RTs is a commonly used method [19], non-autonomous LTR-RTs often lack certain protein domains [20], making it challenging to unify them into a single tree for clustering. LTR_Stream overcomes this limitation. Additionally, the modules identified by LTR_Stream can be used to identify and differentiate fragmented LTR-RTs.

Using LTR_Stream, we subdivided the most abundant Retand LTR lineage in three *Papaver* genomes into 20 sub-lineages. At this resolution, we systematically compared the dynamic changes of LTR-RTs across different subgenomes. For instance, we identified molecular genetic markers that provide evidence of Retand elements spreading between species in a retrovirus-like manner. By leveraging modules identified by LTR_Stream, we detected fragmented LTR-RTs in the whole genome, uncovering their activity diversity across subgenomes and their genomic impacts, such as influencing TAD-like structures. Our analysis of Papaver species using LTR_Stream provides a reference case for studying the interactions between LTR-RTs and host genomes in non-model organisms. We believe LTR_Stream has significant potential to uncover previously unknown evolutionary mechanisms of LTR-RTs in other non-model species.

## RESULTS

### Overview of LTR_Stream

LTR_Stream is designed to cluster LTR-RTs at the sub-lineage level, providing a more nuanced view compared to traditional methods (Fig. 1A). It takes as input the nucleotide sequences of intact LTR-RTs (those containing two identical or very similar LTRs) belonging to the same LTR-lineage. While it can theoretically handle a mix of LTR-RTs from different LTR-lineages, using elements from a single LTR-lineage is recommended for optimal results. LTR_Stream initially performs self-blast with these LTR-RTs, using the results to construct a homology graph. By detecting the connectivity of the graph, LTR_Stream segments each LTR-RT into distinct modules (Figure S1, see methods), which represent potentially functional or structural conserved regions shared among part of LTR-RTs. LTR_Stream then simplifies the representation of a LTR-RT as a sequential order of modules (module sequence) that reflects the order and composition of LTR-RTs. To cluster LTR-RTs, LTR_Stream calculates a distance matrix by estimating pairwise Levenshtein distances [21] between module sequences and then reduces it to a 3D space. Finally, a self-designed layer-by-layer clustering algorithm identifies sub-lineages of LTR-RTs within this space (see methods).

**Figure 1:**
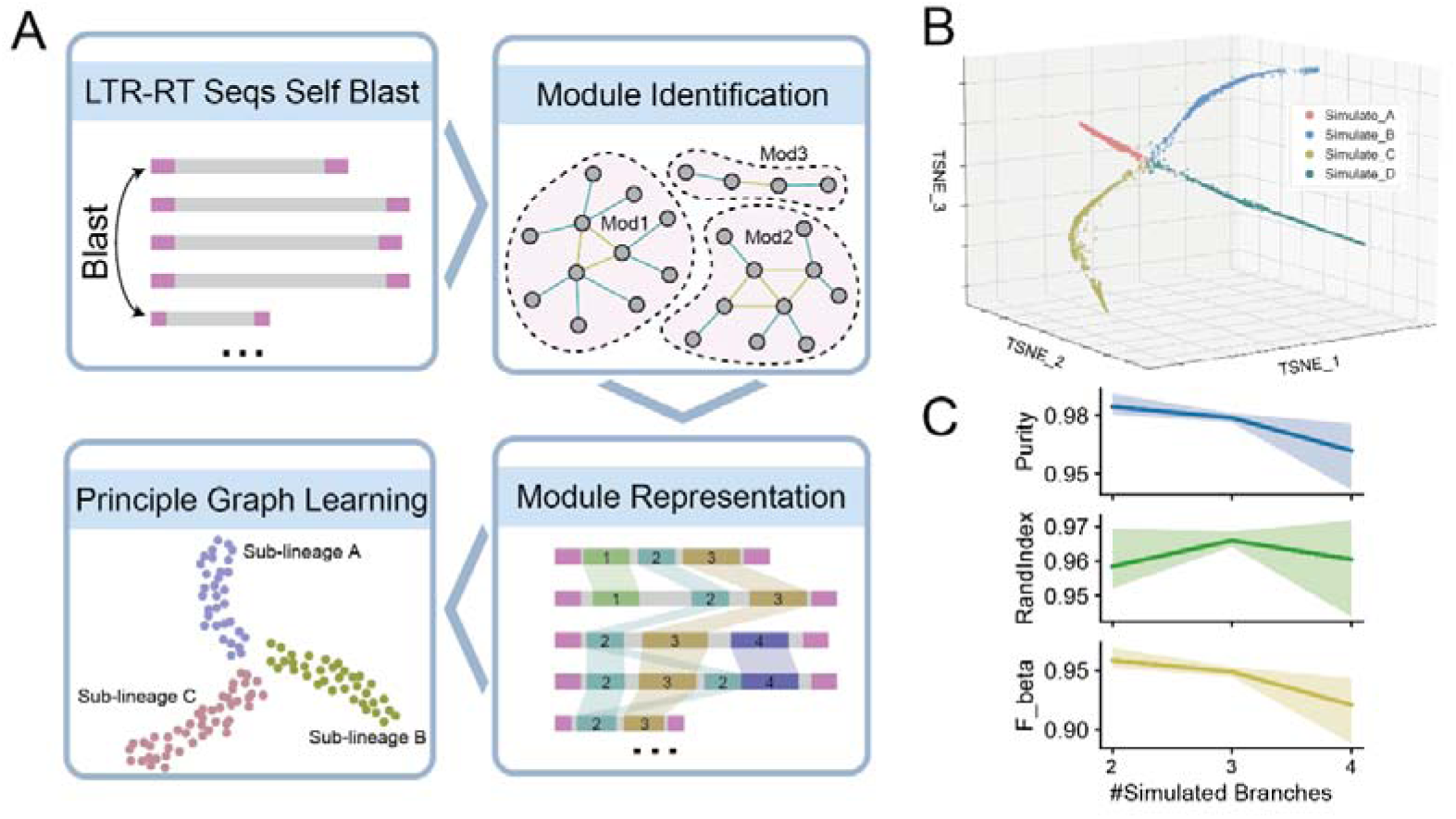
Workflow of LTR_Stream and its performance on simulated datasets. **A.** Flow chart showing the four main procedures of LTR_Stream. **B.** 3D dot plot showing the reconstructed trajectories of LTR-RTs on one simulated dataset with four simulated paths. Each dot represents one module sequence that extracted from one or several LTR-RTs. Colors of dots indicate original simulated path ID. **C.** Line plots showing clustering performance on nine simulated datasets.

### Evaluation of LTR_Stream on simulated and real-world datasets

To assess LTR_Stream’s ability to accurately cluster LTR-RTs at the sub-lineage level, we first tested it on nine sets of simulated data (Table S1). We began by random selection of three real LTR-RT sequences (Ale, CRM, and Tork) from the *Gossypium herbaceum* genome [14]. Each of these sequences were considered the starting point (ancestor) for each simulated dataset. Next, we created simulated independent evolutionary paths for these LTR-RTs, mimicking the four types of changes (bursts, indels, structural variations, and elimination) (Fig. S2). LTR_Stream successfully distinguished the different simulated evolutionary paths (Fig. 1B, S3-5). Various metrics commonly used to evaluate clustering performance (purity, Rand index and F beta score) confirmed LTR_Stream’s effectiveness in correctly clustering simulated LTR-RTs (Fig. 1C).

Having confirmed LTR_Stream’s effectiveness with simulation, we next evaluate LTR_Stream on real LTR-RTs from two closely related plant groups (the *Papaver* group and *Gossypium* group, Table S2). We classified LTR-RTs from each of these two closely groups into LTR-lineages (Table S3-4) and selected six of them (Retand, Ale, CRM, Ogre and Tork of the *Papaver* group, Tekay of the *Gossypium* group) as datasets for evaluation. Notably, the six datasets contain varying numbers of LTR-RTs, ranging from over 10,000 to a few hundred, allowing a comprehensive evaluation of LTR_Stream across different data scales. The results are presented in Fig. 2 and Fig. S6-10, respectively. For example, Fig. 2A illustrated sub-lineage clustering (labeled from A to T) of the Retand set at different subviews. To evaluate these sub-lineages, we randomly selected twenty LTR-RT sequences from each and calculated their pairwise sequence distances (Fig. 2B). We found that in the most of cases (51 of 53), sequences from different sub-lineages were significantly more divergent than these from the same sub-lineage (with fold changes exceeding one in Fig. 2C, Fig. S6-10B). This indicates a higher degree of nucleotide sequence similarity within each sub-lineage. In the case of Retand LTR-RTs from the *Papaver* group, to further validate sub-lineage clustering of LTR_Stream, we utilized their conserved protein domains to construct phylogenetic trees [19,22]. Given the significant presence of nonautonomous LTR-RTs lacking conserved protein domains (Fig. S11), we first constructed an evolutionary tree using the most widely distributed PROT domain across various sub-lineages. The tree confirmed most (16/19) of the sub-lineages (Fig. 2D). For the sub-lineages R, T, and S that exhibit clustering confusion on the tree, we found significant differences in their nucleotide sequences (Fig. S12), validating the rationale behind LTR_Stream clustering. To validate the clustering of sub-lineage M, which only possesses the GAG domain, we also constructed an evolutionary tree using this domain (Fig. S13A). The tree showed that sub-lineage M clusters with sub-lineage L, with the main difference being that L contains additional PROT domain (Fig. S13B). In summary, these analyses confirm the reliability of LTR_Stream and show that nucleotide sequence clustering with LTR_Stream can achieve higher resolution than clustering based on conserved protein domains. Additionally, the estimated insertion time of different sub-lineages varied, suggesting the reliability of the results (Fig. 2C, Fig. S6-10C).

**Figure 2:**
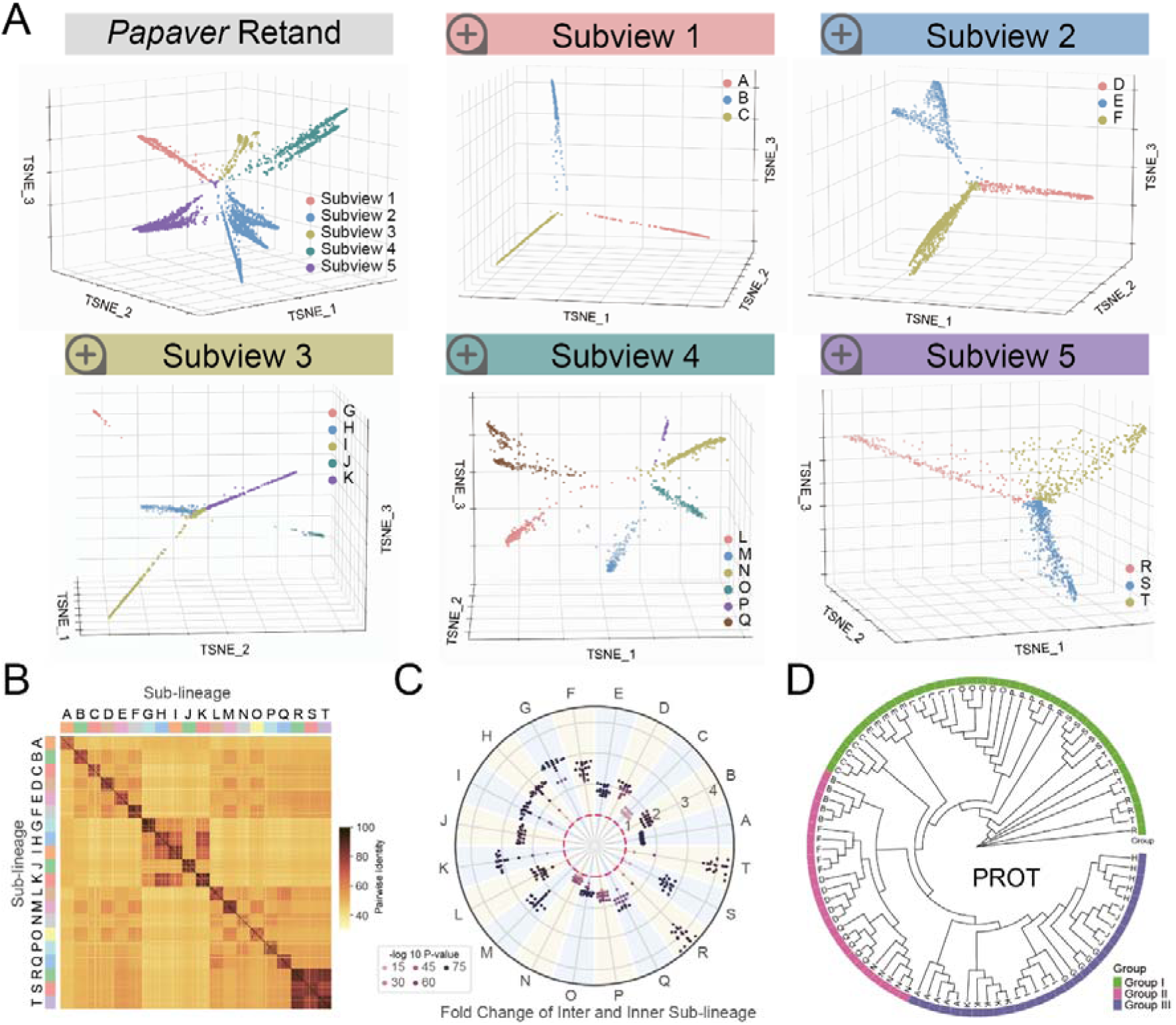
Sub-lineage clustering by LTR_Stream on Retand LTR-RTs of the *Papaver* group. **A.** 3D dot plots showing different sub-lineages in different subviews. Each dot represents one module sequence. Colors of dots indicate different subviews or sub-lineages. **B**. Heatmap showing pairwise identity of randomly selected LTR-RTs. **C.** Swarm plot showing fold change of inter and internal sub-lineage pairwise sequence distances. Colors of dots indicate P-value by Wilcoxon rank-sum test. The red dashed circle represents a fold change of one. **D.** Rootless phylogenetic tree showing randomly sampled PROT sequences from the nineteen sub-lineages (without sub-lineage M). The outermost color blocks represent different Groups.

### Sub-lineage level LTR-RT clustering reveals differential LTR-RT distributions between subgenomes

The clustering of LTR-RTs at a finer sub-lineage level using LTR_Stream provides a novel perspective for studying and comparing those closely related genomes/subgenomes. We performed an in-depth analysis of the distribution of Retand LTR-RTs across different genomes/subgenomes in three *Papaver* species (*P. rhoeas, P. somniferum* and *P. setigerum*) [13]. As shown in Fig. 3A, these three species underwent complex differentiation and allopolyploidization events. Historically, five subgenomes (SG1-4 and PRH) have been identified [23], with PRH diverging early (∼7.1 Mya) from the other four subgenomes and evolving independently into the later *P. rhoeas*. In contrast, SG1 and SG2, as well as SG3 and SG4, merged after their differentiation, leading to new species, with the former being the ancestor of *P. somniferum*, and the latter not yet identified in existing species. These two species, each containing two subgenomes, later merged and evolved into *P. setigerum*. Therefore, *P. rhoeas*, *P. somniferum*, and *P. setigerum* each contain one, two, and four subgenomes, respectively. Due to the relatively short divergence time (∼0.66 Mya), SG1 and SG2 in *P. setigerum* are highly similar to those in *P. somniferum*.

**Figure 3:**
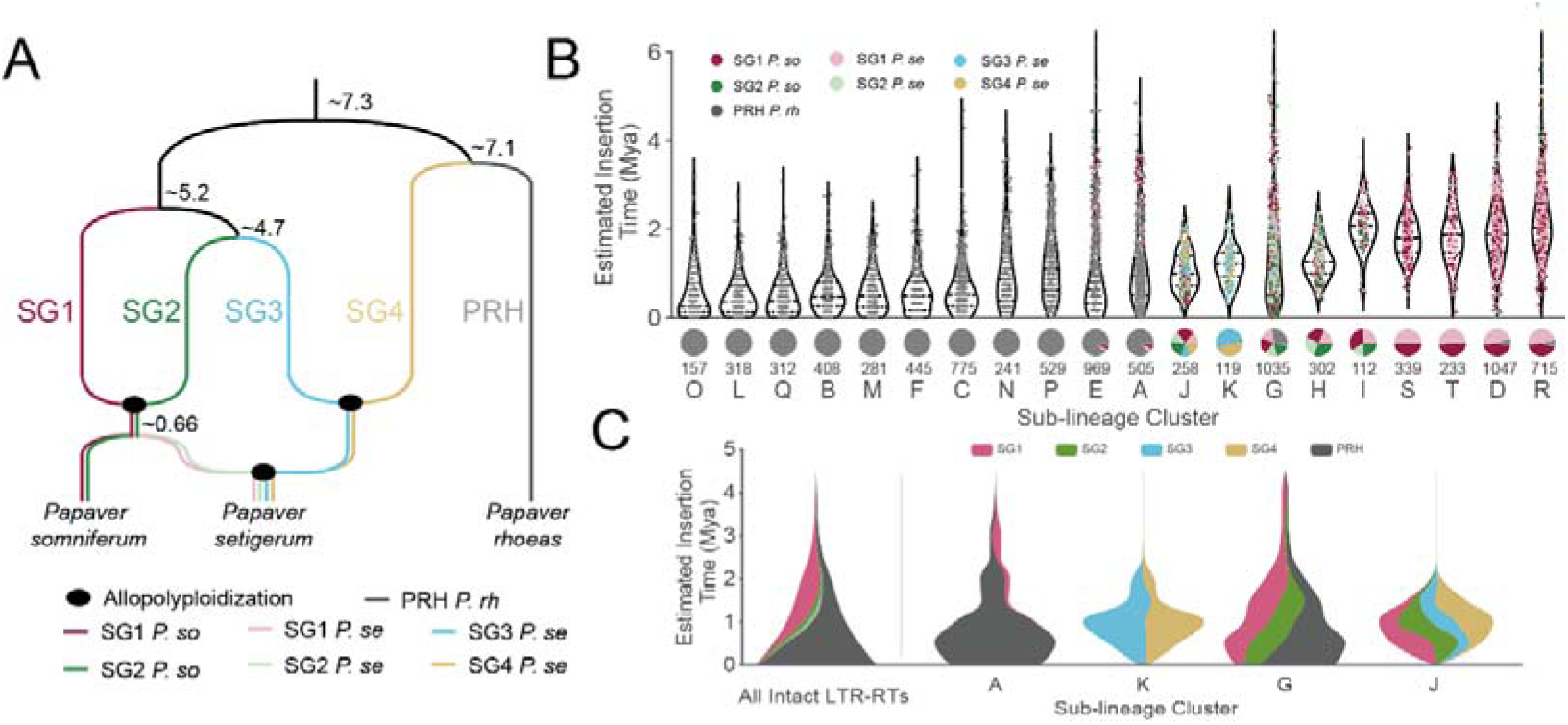
The distribution of 20 sub-lineages of Retand LTR-RTs across the five *Papaver* subgenomes. **A.** Species phylogenetic tree of three *Papaver* species. The divergence times (Mya) of subgenomes are indicated in the figure (not to scale). Black ellipses represent allopolyploidization events. Lines in different colors represent different subgenomes. **B.** Top: Violin plot showing the distributions of the estimated insertion time for each sub-lineage. The color of the dots represents different subgenomes. Bottom: The pie chart illustrates the proportions of different sub-lineages in each subgenome. The number of intact LTR-RTs of each sub-lineage is indicated below pie charts. **C.** The theme river plot illustrates the dynamic changes of LTR-RTs over different time periods across the five subgenomes. The far-left panel shows the overall dynamic changes of LTR-RTs across the five subgenomes, while the four panels on the right display the dynamic changes of LTR-RTs in each of the four sub-lineages.

We found that LTR_Stream effectively reveals the differential distribution of LTR-RTs across the five subgenomes. As shown in the pie charts of Fig. 3B, we observed that 11 sub-lineages are predominantly distributed in the PRH, four are mainly found in SG1 (with similar quantities in SG1 *P. se* and SG1 *P. so*), and five are shared across more than one subgenome. Analysis of insertion times indicates that sub-lineages predominantly located in PRH have the most recent insertion times, suggesting that LTR-RTs in PRH are currently more active than those in the other four subgenomes. Fig. 3C illustrates the evolutionary patterns of sub-lineages across different subgenomes. For example, sub-lineage A was present in both PRH and SG1 before 2 Mya, but it subsequently underwent an expansion in PRH while nearly disappearing in SG1. At the same time, we observed sub-lineages, such as K, G and J shown in Fig. 3C, that expanded almost simultaneously in multiple subgenomes. This reflects the differing adaptive strategies of these sub-lineages in their long-standing evolutionary arms race [24] with the host genomes.

### Sub-lineage level LTR-RT clustering demonstrates that autonomous Retand LTR-RTs could spread between species like retroviruses

The sub-lineages clustered by LTR_Stream in different subgenomes can serve as molecular markers for host genome evolution and also provide insights into how LTR-RTs spread and evolve across the three *Papaver* species. Based on the proportion of conserved protein domains, we classified these sub-lineages into three groups (Fig. 4A). Sub-lineages of Group I mainly contain GAG and PROT domain, those of Group II lack GAG domain, and mainly of Group III include all six domains. For Group II, to rule out potential annotation loss due to divergence between GAG sequences and the database, we confirmed the distance from the 5’ LTR to the start of the PROT ORF and found that this distance is significantly shorter in Group II sub-lineages compared to the other two groups (Fig. S14). Groups I and II, lacking key protein domains, are non-autonomous LTR-RTs, while most in Group III are autonomous LTR-RTs. The phylogenetic trees constructed based on the consensus sequences of GAG and PROT (Table S5) for each sub-lineage suggest that sub-lineages in Group I likely shared a protein domain loss event (Group II as well; Fig. S13). Due to the recent divergent time (Fig. 3A) [23] and similar transposon distributions of SG1 *P. so* and SG1 *P. se* (also SG2 *P. so* and SG2 *P. se*) (Fig. 3B), we assume they share most transposon insertions and do not distinguish them in this section. The pie chart in Fig. 4A shows the subgenome distribution of these 20 sub-lineages. Sub-lineages with >80% distribution in a specific subgenome are defined as SG-specific sub-lineages (including 15 sub-lineages, annotated with blue line in Fig. 4A), while the remaining five (sub-lineage I, G, H, J and K, annotated with red line in Fig. 4A) are SG-shared sub-lineages. We found all of the five SG-shared sub-lineages belong to Group III, indicating a broader distribution of autonomous LTR-RTs in the three species.

**Figure 4:**
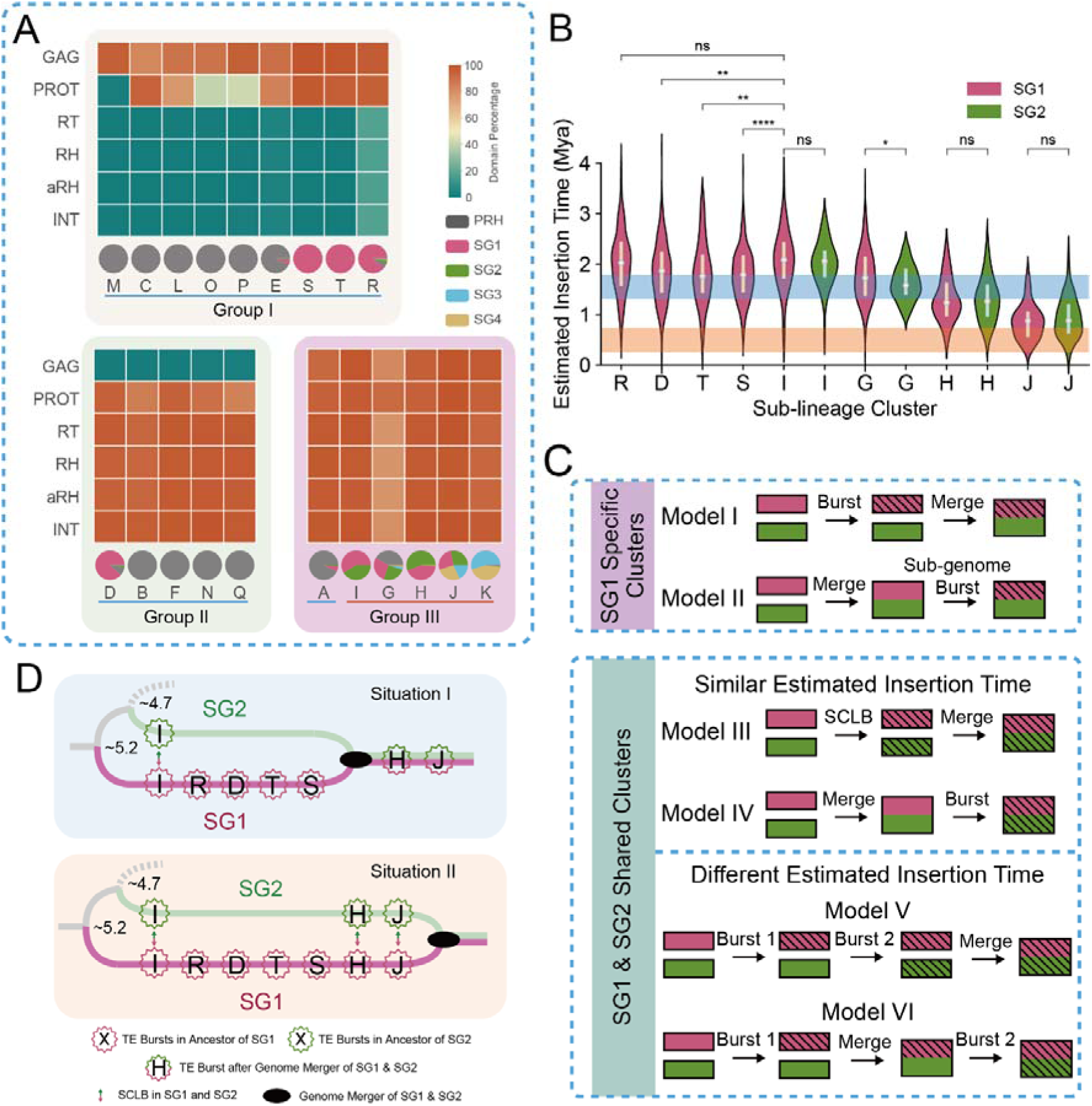
Using sub-lineages as molecular markers to investigate the spread patterns of LTR-RTs in the progenitors of different *Papaver* subgenomes. **A.** Heatmap showing protein domain percentages of the twenty clustered sub-lineages by LTR_Stream. The pie chart at the bottom shows the distribution across different subgenomes (SG1 *P. so* and SG1 *P. se* were merged into SG1, SG2 *P. so* and SG2 *P. se* were merged into SG2). **B.** Violin plots showing estimated insertion time of SG1 and SG2 specific sub-lineages. Colors of violins indicate in which subgenome (SG1 or SG2) LTR-RTs are. The orange shadow denotes the previously reported allopolyploidization time, while the blue shadow denotes another possible time we proposed. Wilcoxon rank-sum test results are also annotated, where ‘ns’ indicates P-value > 0.05, ‘*’ indicates 0.01 < P-value≤0.05, ‘**’ indicates 0.001 < P-value≤0.01, ‘***’ indicates 0.0001 < P-value≤0.001, ‘****’ indicates P-value ≤ 0.0001. **C.**

Both SG-specific and SG-shared sub-lineages are important molecular markers. For those sub-lineages that burst after the sub-genome progenitors diverged from their common ancestor, SG-specific bursts typically accumulate during periods of independent evolution of progenitors of sub-genomes (Model I in Fig. 4C), while the SG-shared sub-lineages may exhibit various scenarios (Model III-VI in Fig. 4C). For those sub-lineages with similar estimated insertion times in two subgenomes, we proposed two possible models:

1. The progenitors of the two subgenomes independently experienced simultaneously cross-species LTR-RT bursts (SCLB) followed by allopolyploidization (Model III in Fig. 4C)
2. The sub-lineages burst occurred after allopolyploidization (Model IV in Fig. 4C)

For those sub-lineages with different insertion time, we also propose two models:

1. The sub-lineages burst in the progenitor of one subgenome and then horizontally transferred to the other (Model V in Fig. 4C)
2. The sub-lineages burst in the progenitor of one subgenome and then, after allopolyploidization occurred, there was another burst across the entire genome (Model VI in Fig. 4C)

Taking the sub-lineages mainly distributed in SG1 and SG2 as an example, we found that sub-lineages T, R, S and D were specific to SG1, while sub-lineages I, G, H and J were shared between SG1 and SG2 (Fig. 4A). The estimated insertion time show that all of these sub-lineages bursts after the progenitors of SG1 and SG2 diverged from their common ancestors (Fig. 4B). For SG-shared sub-lineages, excepting sub-lineage G, sub-lineages I, H, and J exhibited similar estimated insertion time across the two subgenomes (Fig. 4B). Sub-lineage I was distinct in that it burst before either of the SG1-specific sub-lineages such as D, T and S. We considered that if sub-lineage I burst after the allopolyploidization (Model IV), then sub-lineages D, T and S should more closely resemble Model II – however, we thought this is less likely, because it did not conform to the current understanding of retrotransposon mechanisms. Hence, we infer sub-lineage I reflected an SCLB event in progenitors of SG1 and SG2 (Model III). Such SCLB event was intriguing, because it indicated that horizontal transfer of LTR-RTs [25,26] between closely related species and the subsequent expansions in host genome could occur very rapidly, more akin to a retrovirus pandemic. For sub-lineages H and J, it was hard to say whether they followed Model III or IV, because there were no subsequent SG1-or SG2-specific sub-lineage bursts. If H and J followed Model IV, then allopolyploidization was estimated to have occurred 1.30-1.79 Mya (that is, the median estimated insertion time between H and T, shown as the blue shadow in Fig. 4B) and so we propose a possible sub-lineage burst scenario (Fig. 4D). However, if H and J followed Model III, the allopolyploidization event should have occurred later (Fig. 4E), which is consistent with a previous report (the estimated time was 0.26-0.74 Mya, shown as an orange shadow in Fig. 4B) [23]. The latter hypothesis implied that at least three SCLB events (including sub-lineage I, H and J) in the progenitors of SG1 and SG2, very strong evidences proving that Retand could spread between species like retroviruses. Interestingly, sub-lineages I, H, and J all belong to Group III, the autonomous LTR-RT group, suggesting that compared to non-autonomous LTR-RTs, these autonomous elements have a greater potential for cross-species transmission. Regardless of which evolutionary scenario depicted in Fig. 4D or 4E is correct, sub-lineage I in SG1 and SG2 serves as a strong molecular marker for a past SCLB event, suggesting that historically, Retand LTR-RTs could spread across species in a manner similar to retroviruses.

Possible models to explain SG1 specific (top) and SG1&2 shared (bottom) sub-lineage bursts. Two subgenomes are denoted with different colors. The shading represents LTR-RT bursts. **D and E.** Two possible sub-lineage burst situations in SG1 and SG2. The color of the lines represents the corresponding subgenome. Circle denotes LTR-RT sub-lineage bursts. The figure is for illustrative purposes only and does not adhere to a strict scale.

### Modules identified by LTR_Stream reveal the diverse activity of LTR-RTs across host subgenomes

During the long-term arms race between host genomes and LTR-RTs, host genomes employ various mechanisms to eliminate LTR-RTs, thereby generating truncated fragments of LTR-RTs [27]. Compared to intact LTR-RTs with paired LTRs, these truncated fragments are more numerous and widely distributed across the genome. Recent evidence suggests that these truncated fragments can influence genome functions, such as transcriptional regulation [28]. Luckily, modules identified by LTR_Stream during the clustering process, which represent homologous nucleic sequence fragments shared among certain LTR-RTs, facilitate the genome-wide identification and differentiation of these truncated fragments.

From Retand LTR-RTs of three *Papaver* species, we filtered 35 modules that showed differential distributions among sub-lineages and exhibited low reciprocal overlap rates (<0.5) (Fig. 5A). Some modules were sub-lineage specific, such as module b956494 (the sixth row from bottom in Fig. 5A), which was predominantly distributed in sub-lineage M (the third column from left) (Fig. 5A). In contrast, some modules were shared by several sub-lineage. For example, b956 (the first row from top in Fig. 5A) were mainly shared by sub-lineages Q, N, F, and D. These modules intrinsically reflect sub-lineage-specific nucleotide sequences. For instance, Fig. 5B illustrates the positional annotations of two sub-lineage D associated modules, b144725 and b956, within a single LTR-RT sequence. These modules can be used to search for homologous sequences across whole genome. For example, Fig. 5C shows the positional annotation of module b956 identified in *P. so*. The module b956 is predominantly distributed in sub-lineages F, N, Q, and D. The first three sub-lineages are primarily distributed in PRH, whereas sub-lineage D is predominantly distributed in SG1 (Fig. 3B). Consequently, b956 is also observed to be mainly located in SG1 in *P. so* (Fig. 5C). By calculating the proportion of these modules within intact LTR-RTs, we estimated the elimination rate of LTR-RTs in subgenomes. Interestingly, we found that PRH, within which Retand is most active, also exhibits the highest elimination rate (Fig. 5D). Using transcriptome data from six tissues (including petal, fruit, stem, leaf, fineroot and stamen) [13] across the three species, we evaluated the transcriptional activity of these modules. Modules in PRH were consistently found to have the highest expression levels across all six tissues (Fig. 5E). Among these tissues, the stamen, as a reproductive organ, may experience reduced methylation due to epigenetic reprogramming, thereby reactivating some LTR-RTs [29]. However, increased transposon expression was only observed in SG2 (including both SG2 *P. so* and SG2 *P. se*, Fig. 5E), suggesting that different subgenomes exhibit varying capacities to suppress transposons following a reduction in methylation. Retand in SG2, which are more active in reproductive organs, may be more prone to hereditary accumulation. Previous studies have reported that more active transposons often exhibit higher elimination rate [24]. To investigate the correlation between activity and elimination at the module level, we analysed their relationship across all five subgenomes but found no strong positive correlation (Fig. 5F). This indicates that different modules exhibit relatively dynamic changes within each subgenome. For instance, modules with higher expression and lower elimination rates tend to be in an expansion phase compared to others. Since SG1 and SG2 in *P. so* and *P. se* diverged relatively recently (∼ 0.66 Mya) [23], and the latter underwent an additional allopolyploidization (Fig. 3A), these genomes provide an excellent system to study the impact of allopolyploidization on transposon activity. Our findings reveal that most modules across both subgenomes show a general decline in overall transcription levels accompanied by an increase in elimination rates (Fig. 5G). While previous studies have reported that allopolyploidization can induce genome shock, resulting in transposon activation [30–34], the opposite pattern is observed in *P. se*. Next, we compared the expression level of fragmented modules and those located within intact LTR-RTs. Indeed, we would expect to observe a higher expression tendency in those within intact LTR-RTs, as fragmented LTR-RTs have already lost their transposition activity. Surprisingly, in SG1 and SG4, we found that expression of fragmented LTR-RTs of three modules was even higher, which raises questions about the potential effects of high expression of fragmented transposons on their transposition behaviour (Fig. 5H). Overall, modules in the more active PRH subgenome tend to show higher expression of intact LTR-RTs (Fig. 5H), suggesting that the expression of intact LTR-RTs plays a more significant role in transposon activity. For instance, we observed that b144725 is significantly more highly expressed in intact LTR-RTs in PRH, while in SG1, there was no significant difference in expression between fragmented and intact LTR-RTs (Fig. 5I). Module b144725’s associated sub-lineages D and S have an earlier insertion time in SG1, while sub-lineages E and F were inserted later in PRH (Fig. 5I), indicating that they still maintain considerable activity in PRH. Our analysis demonstrates that the modules identified by LTR_Stream provide a more comprehensive understanding of the diversity of LTR-RT activity across different subgenomes, offering a more nuanced perspective for studying the arms race between LTR-RTs and the host genome.

**Figure 5:**
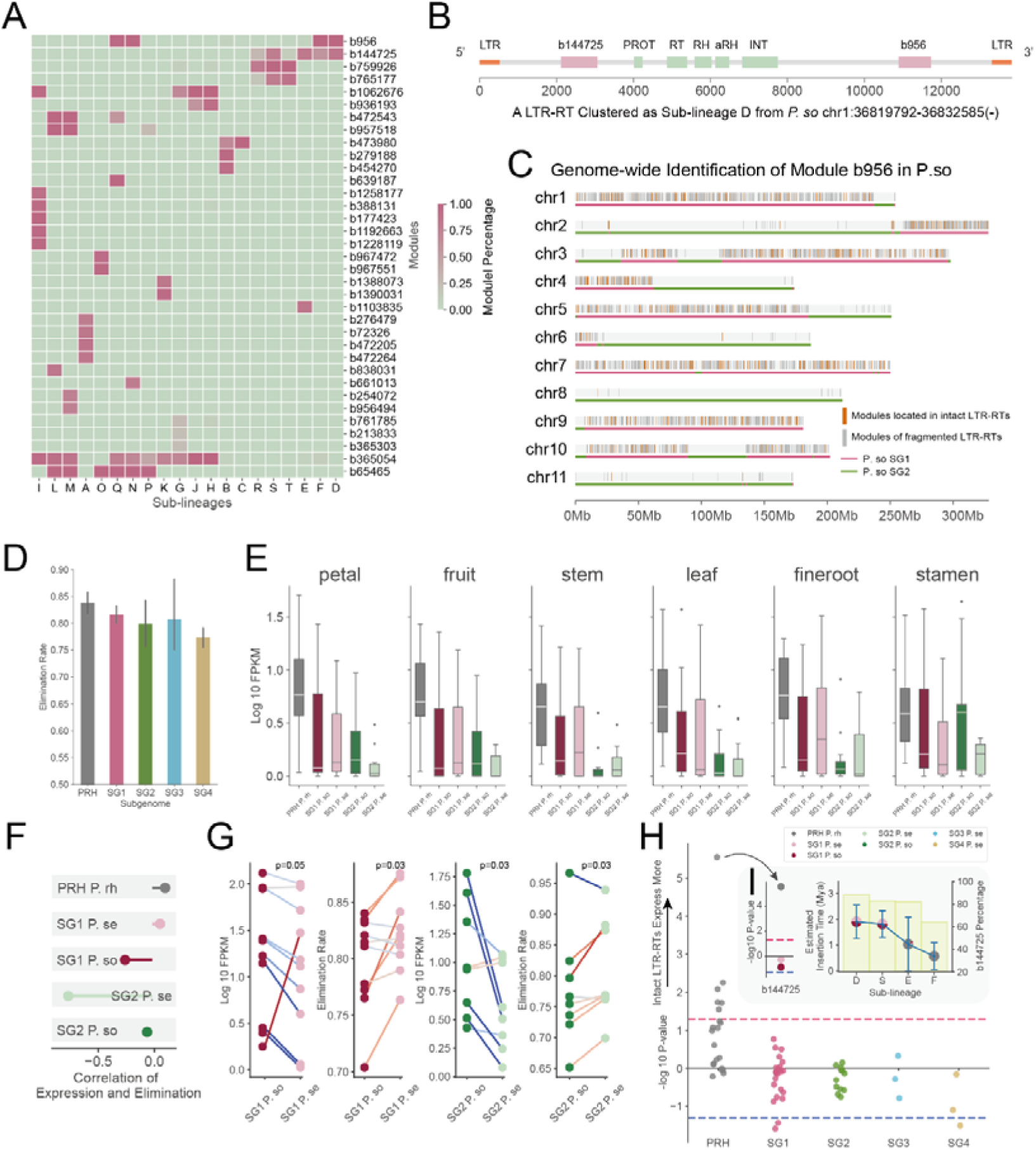
Analyse the activity diversity of Retand LTR-RTs across different *Papaver* subgenomes at the module level. **A.** The heatmap illustrates the distribution of different modules across different sub-lineages. The numbering starting with “b” represents the module ID. **B.** An example shows the position of modules on an LTR-RT sequence. The light green rectangle represents the conserved protein ORF. The pink rectangle represents the modules identified by LTR_Stream. **C.** Using module b956 as an example, its distribution on the *P. so* genome is shown. The modules located in the intact LTR-RT regions are marked in orange, while those located in truncated fragments are highlighted in gray. The two subgenomes are indicated by lines in different colors. **D.** The bar chart displays the elimination rate of LTR-RTs in different subgenomes, assessed using modules. **E.** The box plot shows the expression levels of modules in PRH, SG1 *P. so*, SG1 *P. se*, SG2 *P. so* and SG2 *P. se* across six tissues. **F.** The lollipop chart illustrates the correlation between module expression levels and elimination rates in different subgenomes. **G.** The dumbbell plot shows the changes in expression levels and elimination rates of Retand LTR-RTs in before and after the last allopolyploidization in *P. se*. **H.** The swarm plot shows the expression level differences between intact LTR-RTs and truncated fragments across the five subgenomes. The red dashed line represents a higher expression in intact LTR-RTs with a P-value of 0.05 (Wilcoxon rank-sum test). The blue dashed line represents a higher expression in truncated fragments with a P-value of 0.05 (Wilcoxon rank-sum test). **I.** Using module b144725 as an example, the plot shows its preferential expression between intact LTR-RTs and truncated fragments across different subgenomes, alongside the activity level of the corresponding sub-lineage in the respective subgenomes.

### Modules identified by LTR_Stream influence nearby TAD-like structure boundaries

Next, we explored the functional impact of the modules identified by LTR_Stream on the genome. Previous studies in wheat have reported that LTR-RTs are involved in higher-order chromatin organization [35]. Using Hi-C sequencing data, we analysed the potential associations between these modules and higher-order chromatin organization across the genome. We discovered that certain modules are closer to TAD-like boundaries compared to random regions (Fig. 6A), such as module b1062676 in SG2 of *P. so*, which we define as near-TAD modules. We propose two possible explanations for this phenomenon: (1) these near-TAD modules contribute to the generation of nearby TAD-like boundaries, or (2) LTR-RTs carrying these near-TAD modules are preferentially inserted near TAD-like boundaries. To determine which of these scenarios better explains this intriguing observation, we used colinear synteny gene pairs [36] across different subgenomes as a framework to identify homologous region pairs (Fig. 6B, 6C). Due to the extensive transposon bursts and accumulation in PRH, we excluded it from this analysis. In detail, we first selected homologous region pairs that contain one Region A (without any near-TAD modules) and one Region B (contains near-TAD modules) (Fig. 6C). We hypothesized that the TAD-like structures in Region A are not influenced by LTR-RTs and therefore represent a state closer to the ancestor compared to Region B. We then compared the TAD-like boundary density between these region pairs. Fig. 6D shows an example of module b1062676 in SG2 *P. so*, that is after TE insertion, interval length of TAD-like boundary of corresponding genome region became shorter, indicating higher TAD-like boundary density. In general, we found that, except for SG1 in *P. se*, the Region A associated TAD-like boundary density was significantly lower than that of Region B across all other subgenomes (Fig. 6E). Moreover, the significance of this reduce was strongly positively correlated with the significance of modules closer to TAD-like boundaries, with correlation coefficients exceeding 0.95 in SG2, SG3 and SG4 (Fig. 6E). These findings support the first hypothesis: LTR-RT insertions into subgenomes indeed influence the nearby TAD-like structures. Next, we selected region pairs unaffected by any LTR-RT insertions (Regions C and D in Fig. 6C) as negative controls for Region A. If the second hypothesis were correct, we would expect the Region A associated TAD-like boundary density to be significantly higher than in Regions C and D. However, as shown in Fig. 6F, this was not the case. On the contrary, we observed that the TAD-like boundary density near Region A was even lower in certain cases, such as those involving modules in SG1 and SG3 (Fig. 6F). Additionally, the significance does not show positively correlated with the significance of modules closer to TAD-like boundaries (Fig. 6F), further disproving the second hypothesis. Thus, we conclude that certain near-TAD modules indeed induce changes in TAD-like structures. Fig. 6H illustrates an example: in homologous pair regions across SG3 and SG2, the insertion of the LTR-RT module b1062676 in SG2 may contribute to the formation of a new TAD-like boundary nearby (indicated by the red dashed line), which is absent in SG3. Notably, compared to SG1 and SG2 in *P. se* (light red and light green in Fig. 6A), the proximity significance of modules to TAD-like boundaries was higher in *P. so* (dark red and dark green). This observation suggests that allopolyploidization events may attenuate the impact of modules on TAD-like boundary formation. Previous studies in plants have shown that the bZIP and TCP transcription factor families can influence three-dimensional chromatin structures and their associated motif are enriched near TAD-like boundaries [37,38]. Consistently, we identified motifs associated with these transcription factors enriched within near-TAD modules (Fig. 6G, Supplementary File S1). Given that these motifs are also enriched near TAD-like boundaries in the corresponding subgenomes (Supplementary File S2), we propose that the presence of TCP and bZIP-associated motifs in these modules is likely the mechanism by which they influence TAD-like structural organization. We observed that modules with higher proximity significance to TAD-like boundaries also shared more TCP- and bZIP-related motifs with each other (Fig. S15). Transcriptomic analysis revealed that b1062676 in SG1 and SG2 maintains a certain level of activity (FPKM around 10) (Fig. 6I), indicating its potential to continuously influence the higher-order 3D structure of its host genome. Based on these findings, we propose a model (Fig. 6J), that is, modules such as b1062676, which carry bZIP and TCP-related motifs, may mediate the formation of new TAD-like boundaries in their vicinity.

**Figure 6:**
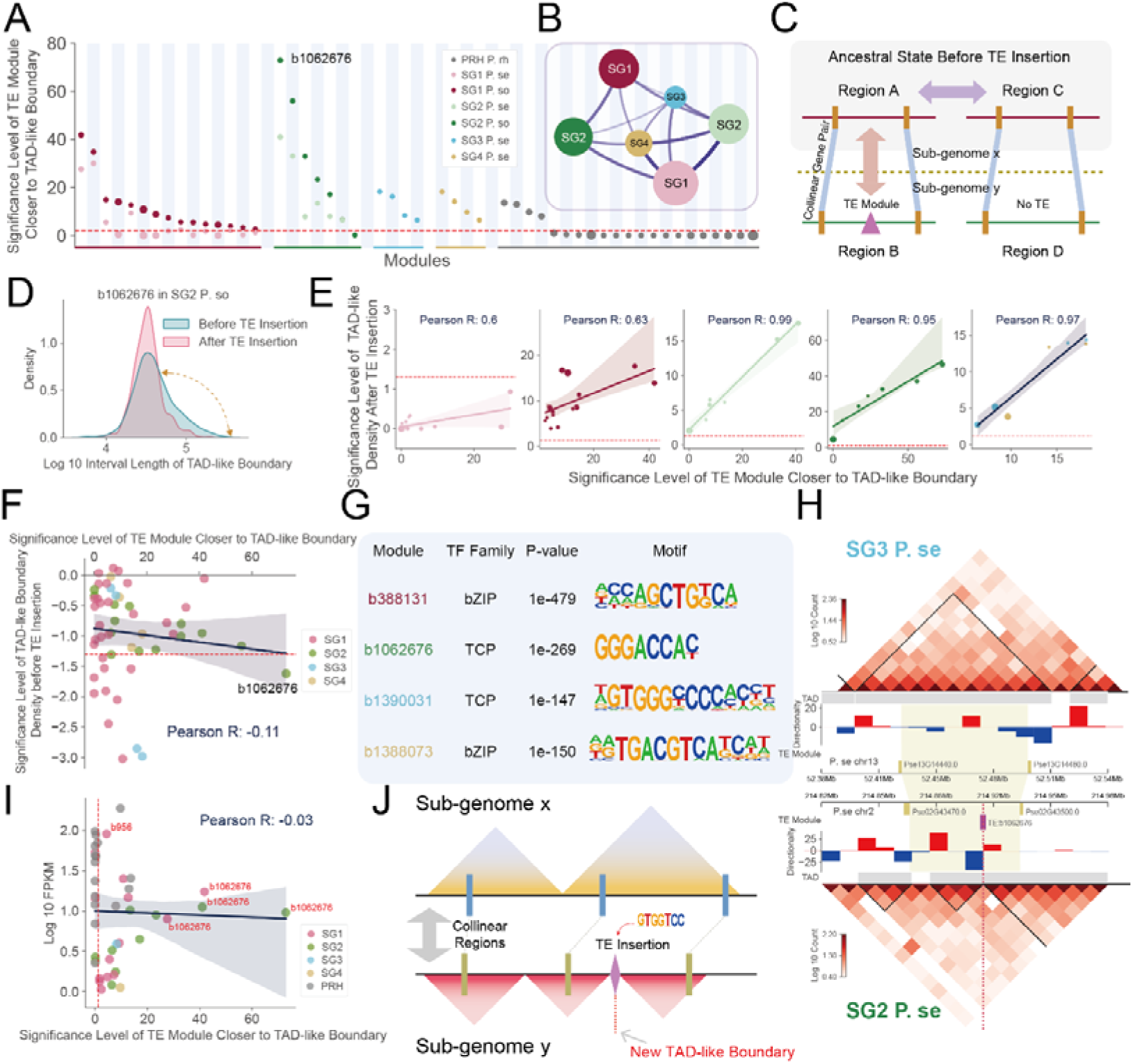
Modules carrying certain TCP or bZIP transcription factor related motifs may mediate the formation of new TAD-like boundaries. **A.** The dot plot shows the significance of proximity between modules and TAD-like boundaries (Wilcoxon rank-sum test). The color of the dots represents different subgenomes. **B.** The graph shows subgenomes that used to identify colinear synteny gene pairs. The number of colinear synteny gene pairs is represented by the width of the edges. **C.** The schematic diagram illustrates how to compare changes in TAD-like structures after LTR-RT insertion. **D.** The density plot shows changes in adjacent TAD-like boundary intervals around module b1062676 after LTR-RT insertion. **E.** The dot plot shows the association between the significance of TAD-like boundary density changes near different modules after LTR-RT insertion (Y-axis) and the significance of module proximity to TAD-like boundaries (X-axis) across different subgenomes (Wilcoxon rank-sum test). The red dashed line represents a P-value of 0.05 for the significance of 0.05. **F.** The dot plot shows the significance of TAD-like boundary density near sites with LTR-RT insertions but before insertion compared to sites without any LTR-RT insertions, along with the significance of module proximity to TAD-like boundaries (Wilcoxon rank-sum test). The color of the dots represents different subgenomes. **G.** The most enriched TCP and bZIP related motifs in the four modules from different subgenomes. Color of the module ID represents the corresponding subgenome. **H.** An example illustrates the changes in TAD-like structures after LTR-RT insertion in SG2 *P. se* (bottom), compared with that in SG3 *P. se* (top). The heatmap shows Hi-C interactions at a 10 kb resolution. The bar chart shows the Directionality Index. The purple rectangles represent modules. The yellow rectangles represent colinear synteny gene pairs identified by MCScanX. **I.** The dot plot shows module total expression level (Y-axis) and its correlation to the significance of module proximity to TAD-like boundaries (X-axis). The color of the dots represents different subgenomes. **J.** A model schematic illustrates that modules carrying TCP- and bZIP-related motifs might mediate the formation of new TAD-like boundaries.

## DISCUSSION

The diversity and widespread distribution of LTR-RTs in plants make them valuable molecular markers for genomic evolution. For example, as early as a decade ago, researchers used PCR-based TE markers to reveal phylogenetic relationships within and between plant species [39]. In recent years, with advances in sequencing technologies, k-mer analysis based on assembled genome have been employed for LTR-RT marker identification [40,41]. Typically, during independent evolution after ancestral species divergence, subgenome-specific distributed TEs may accumulate, while allopolyploidization events halt this biased accumulation. By comparing subgenome ancestor divergence time and the insertion time of subgenome-specific LTR-RTs, these methods can identify allopolyploidization events and estimate their timing. The high-resolution LTR-RT clustering achieved by LTR_Stream facilitates in-depth analysis of LTR-RT burst and propagation patterns in closely related species. Take three *Papaver* species as an example, we observed that the ancestors of two subgenomes experienced simultaneous bursts of the same sub-lineage during their independent evolution following species divergence, with such events likely occurring three times in the recent four million years. Although horizontal transfer of LTR-RTs among species is not uncommon, the near-simultaneous bursts of same LTR-RT sub-lineages in closely related species was rarely reported. We hypothesize that these simultaneously cross-species LTR-RT bursts (SCLBs) suggest that these sub-lineages may have once possessed the ability to spread rapidly between species, akin to plant viruses. However, the precise mechanisms underlying the cross-species transmission of Retand transposons remain unclear. On one hand, Retand lacks the ENV protein, which is essential for interacting with cell surface receptors and mediating fusion. On the other hand, the plant cell wall poses a physical barrier to intercellular LTR-RT transmission. We found that all sub-lineages potentially associated with SCLBs are autonomous and identified two additional ORFs encoding transmembrane proteins in Retand (Figure S16), which may functionally substitute for ENV proteins. These findings provide valuable insights for further researches about mechanisms of LTR-RT cross-species transmission. Additionally, further exploration of how host genomes counteract such cross-species transmission threats is warranted, as it may offer critical references for addressing potential plant viral threats in the future.

In addition to serving as molecular markers, the insertion of TEs can directly or indirectly influence plant genome function. For example, numerous studies have reported that TEs carrying specific cis-regulatory motifs can affect the expression of nearby genes, thereby reshaping gene regulatory networks [7,42]. In recent years, the impact of TEs on higher-order chromatin structures has also been increasingly documented in plants [43]. For instance, in cotton, it was reported that, TEs have been linked to transitions between active and inactive chromatin regions, and species-specific TE expansions may mediate the formation of species-specific TAD boundaries [44,45]. However, the specific TEs involved and their underlying mechanisms remain unclear. Leveraging high-resolution clustering results from LTR_Stream, we compared syntenic gene modules and found that the insertion of certain LTR-RT modules correlates with a significant increase in TAD boundary density in their vicinity. Further analysis revealed that these modules are enriched with binding motifs for TCP and bZIP transcription factors. Given that TCP and bZIP motifs are also enriched at TAD boundaries in plant such as rice [37] and cotton [45], we hypothesize that TE-introduced motifs may facilitate the formation of new TAD boundaries in adjacent regions. Although these transcription factors are considered potential regulators of TADs in plants, the specific insulators mediating TAD formation remain unidentified, and further research is needed to elucidate the underlying mechanisms.

TEs and polyploidization are considered two major driving forces in plant genome evolution [24]. Previously, some hypotheses and evidence suggested that the genomic shock induced by polyploidization could trigger massive TE expansions [33,46]. However, recent studies have shown that polyploidization does not necessarily lead to TE bursts in many species [33,46]. *P. setigerum*, formed through the allopolyploidization of progenitor of *P. somniferum* and an unknown species, provides an ideal system to study TE dynamics under polyploidization. Using high-resolution clustering results from LTR_Stream, we found that, compared to *P. somniferum*, the potential threat of Retand to the genome in *P. setigerum* is significantly reduced, evidenced by lower expression levels and higher elimination rates. Notably, we observed a weakened association between TE modules and TAD boundaries after allopolyploidization, although the mechanisms driving this change remain unclear. We noticed that previous studies have shown that polyploidization itself can alter higher-order chromatin structures, such as A/B compartment transitions and TAD reorganization [43]. For example, in soybean, changes in epigenetic modifications (e.g., H3K4me3, H3K9me2, and CG/CHG methylation levels) during polyploidization may drive TAD restructuring [47]. This mechanism parallels findings in animal genomes, where epigenetic changes leading to gains or losses of local transcriptional activity can promote the formation or loss of TAD boundaries [47]. Our study demonstrates that the impact of TEs on the genome is attenuated in *P. setigerum* following polyploidization. Furthermore, considering the significantly higher TE activity in P. rhoeas and *P. bracteatum* [48], which have not undergone recent polyploidization, we propose that polyploidization may serve as an adaptive strategy in *Papaver* species to counteract TE expansion. Given that TE loss has also been reported in other polyploid species [33,46], we further speculate that this mechanism may be a widespread phenomenon in plants, which needs further investigation.

## METHODS

### LTR-RT representation with module sequence

LTR_Stream clusters LTR-RT DNA sequences by first performing self-BLAST with BLASTn (v2.11.0+) [49]. This comparison helps identify homologous regions within the sequences. LTR_Stream then constructs an undirected graph model based on these comparisons. LTR_Stream segments LTR-RT sequences into different sections and identifies them as distinct modules, representing each original LTR-RT sequence as a module sequence. Specifically, for a given self-BLAST result, LTR_Stream builds an undirected graph *G*. The vertices of *G* represent sections of LTR-RT sequences that are identified homologous to other sequences. For any pair of vertices *v* and *j* in the graph, we add an edge *(v, j)* to *G* if either of the following conditions is met (Fig. S1):

1. Section *v* and *j* are identified by BLAST as a homologous pair, and e-value is less than a given threshold.
2. Section *v* and *j* belong to a same LTR-RT, and the mutual overlap of their positions exceeds a given threshold.

Since LTR-RTs often share a lot of similarity, BLAST typically provides a large number of homologous alignment records. To manage this complexity, LTR_Stream limits the total number of alignment records by setting the maximum output records for each LTR-RT. LTR_Stream then efficiently counts the connected components in large graphs using a disjoint-set. Vertices belonging to each connected component are then merged into a module (Fig. S1). According to the original positions of each module, LTR_Stream represents each LTR-RT as a module sequence. In cases where there are too many modules, LTR_Stream represents only those LTR-RTs with the most widely distributed modules.

### Sub-lineage level LTR-RT clustering

LTR_Stream takes the module sequences generated from the previous step and calculates Levenshtein distances [21] between different module sequences and reduces the distance matrix to a 3D space using t-SNE [50]. Due to accumulation of mutations, we expect that older LTR-RTs would have fewer identifiable homologous segments compared to more recently-inserted LTR-RTs. Consequently, we assume that older LTR-RTs would have shorter module sequences and relatively smaller Levenshtein distances when compared with each other. Hence, older LTR-RTs are expected to cluster closer together near the center, while more recent ones, from different evolutionary branches, would be positioned further out in the 3D space. Next, LTR_Stream calculates the angular differences between different module sequences relative to the center, constructing a new distance matrix. LTR_Stream clusters these with Agglomerative Clustering [51], optimizing the clustering with ElPiGraph [52] to reconstruct modular sequence trajectories. Part of the code framework references STREAM [53]. In some cases, LTR-RTs might have complex evolutionary histories, making it difficult to group them definitely. For such cases, LTR_Stream focuses on specific category and refines the clustering iteratively until either the dimension reduction no longer fits the assumption or it reaches the manually set maximum iteration depth. This strategy resembles hierarchical clustering, but due to the sparsity of the features extracted by LTR_Stream, we do not guarantee the intermediate hierarchical clustering. To evaluate the final clusters (sub-lineages), LTR_Stream randomly selects several LTR-RT sequences from each cluster, calculates their pairwise identities, and employs the Wilcoxon rank-sum test to assess whether intra-class similarity significantly exceeds inter-class identity. This helps to adjust parameters and refine the clustering if necessary. Finally, LTR_Stream consolidates the final clustering results obtained from different perspectives and saves them in a TSV (Tab-Separated Value) format file. LTR_Stream also outputs the coordinates of the module sequences in the final view, facilitating the filtering-out of the lower-confidence samples near the origin for users.

### Data simulation

For a given LTR-RT sequence, we simulate LTR-RT bursts iteratively, introducing random structural mutations (including insertions, deletions, and inversions) at each iteration to construct an evolutionary trajectory. To faithfully simulate the evolution of LTR-RT under natural conditions, we reference the model proposed by Drost, H. G. [54], introducing indels that occur along host genome as well as LTR-RT elimination. To test LTR_Stream, we randomly selected one sequence from each of the three LTR-lineages in the *Gossypium herbaceum* genome (Ale, CRM, and Tork) as the ancestral sequences, simulating two to four independent evolutionary paths originating from them.

### Genome wide LTR-RT identification from closely related species

We downloaded the genome assemblies of three *Papaver* [13] and four *Gossypium* species [14,45], as detailed in Table S2. Then, we ran LTR_FINDER (v1.2) [55] with default parameters and LTRharvest (v1.6.2) [56] with parameters “-minlenltr 100 -maxlenltr 7000 - mintsd 4 -maxtsd 6 -motif TGCA -motifmis 1 -similar 85 -vic 10 -seed 20 -seqids yes” on these assemblies. Further, LTR_retriever (v2.9.0) [57] was used to filter high-quality LTR-RTs and estimate insertion times of LTR-RTs. Nucleotide sequences of LTR-RTs were extracted by BEDTools (v2.30.0) [58], which were then used as input to LTR_Stream. The insertion times of each LTR-RT were estimated by LTR_retriever with synonymous substitutions per site per year considered to be 8.1□×□10^−9^ for the *Papaver* species [13] and 3.5 × 10^−9^ for the *Gossypium* species [14].

### LTR-RT protein domain analysis

LTR-RT protein domains were annotated by DANTE (v1.0.0) [59], using the database viridiplantae_v3.0, downloaded from the REXdb [19]. For, evaluationg sub-lineage clustering of LTR_Stream, five amino acid sequences of PROT and GAG were randomly selected from each cluster (50% LTR-RTs near origin will be filtered out). Then, we ran MUSCLE [60] with default parameters on these sequences and constructed phylogenetic trees with FastTree (v2.1.11) [61]. Ggtree (v3.10.0) [62] was used to plot and annotate the two phylogenetic trees.

To clarify the phylogenetic relationships between sub-lineages, we randomly selected 50 GAG and PROT protein sequences from each sub-lineage to construct consensus domain sequences. Then, for each protein in each sub-lineages, we ran MUSCLE [60] with default parameters and generate consensus sequence with biopython(v1.81). Then, with a amino acid sequence from Athila lineage as the outgroup, we used MUSCLE, FastTree and ggtree to generate phylogenetic tree (as mentioned previously).

### Genome-wide module identification and expression quantification

Module with diffScore more than 0.8 was selected, where diffScore was defined as range of module percentages of different clusters. For each module, we randomly selected ten sequences as seeds for searching genome wide homologous regions (BLASTn with parameter ‘-evalue 1e-5 -outfmt 6 -max_target_seqs 10000’). We retained only homologous regions that are at least 50% of the seed length. Considering that homologous regions identified by different seeds may overlap, we merged these regions and assigned the median of their start and end points to the merged region. Next, considering the potential overlap between modules, we filtered out modules that overlapped with longer modules to minimize redundancy (defined as having a median overlap greater than 0.2). The coordinate information of these non-redundant modules is exported to a GTF file.

For module expression quantification, we first used STAR (v2.7.8a) [63] to align RNA-seq data from six tissues (including petal, fruit, stem, leaf, fineroot and stamen) [13] across three *Papaver* species (including *P. rhoeas, P. somniferum* and *P. setigerum*) to their respective genomes [13]. The parameters for STAR were set as ‘--outSAMtype BAM SortedByCoordinate --sjdbOverhang 149 --readFilesCommand zcat -- outFilterMultimapNma× 1000 --winAnchorMultimapNma× 1000’, according to the recommendations of TEtranscripts [64], aiming to maximize the alignment of reads to repetitive regions. Then, TECount (v2.2.3) [64] was used to count of each module and FPKM was estimated as [65]:

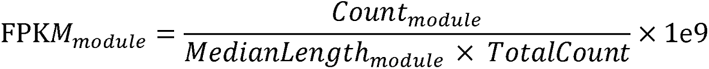

### 3D genome analysis

Hi-C data of *P. somniferum* and *P. setigerum* were downloaded from the NCBI Sequence Read Archive (SRA) database with accession PRJNA720042 [13]. Juicer (v1.5.7) [66] with parameter ‘-s DpnII’ was used for counting interactions. TAD-like structure was called by DI method [67]. For each module, we calculated distances of homologous regions to the closest TAD-like boundary and assessed significance against randomly generated regions (of the same number and with the same length) using the Wilcoxon rank-sum test. Colinear synteny gene pairs was identified by MCScanX [36] across different subgenomes. TCP and bZIP motifs were downloaded from the PlantTFDB database [68].Motif enrichment was performed by HOMER [69]. For each module, enrichment was performed by comparing module sequences to 3000 randomly selected regions (with median length of module sequences) of the corresponding subgenome. For TAD-like boundary, enrichment was performed by comparing flanking 10kb of the TAD-like boundary to 5000 randomly selected regions of the corresponding subgenome.

### Parameter description and adjustment guideline

User-specific datasets require appropriate parameter adjustments to achieve optimal clustering results. LTR_Stream’s parameters are divided into three main categories. The first category includes *minOverlap* and *topModNum*, used for identifying and selecting modules. The second category involves t-SNE-related parameters for dimensionality reduction, such as perplexity, earlyExaggeration, and learningRate. The third category includes parameters used in ElPiGraph. Among these, *minOverlap*, *topModNum*, and perplexity are particularly sensitive to the dataset and significantly impact clustering results. To handle this, LTR_Stream provides visualizations of intermediate results to assist users in tuning these three parameters.

*minOverlap* controls the merging of nearby modules, with a smaller value leading to more merging. *topModNum* defines the number of most frequent modules selected. Before clustering, LTR_Stream provides the percentage coverage of LTR-RTs by different modular sequence length under different *topModNum* settings. A too-small *minOverlap* merges more modules, causing information loss and generally shorter module sequences (Fig. S17A). We recommend that the difference between the coverage at a modular sequence length of 1 near the saturation point and that at length 10 should not exceed twofold (Fig. S17B). A too-large *minOverlap* results in a jagged increase in the coverage curve (Fig. S17C) due to nested modules, potentially introducing outliers. Ideally, the curve should rise as smoothly as possible (Fig. S17B). We recommend setting *minOverlap* between 0.8 and 0.99. For *topModNum*, we suggest a value slightly above the saturation point of coverage for modular sequences of length 2 or 3. If outliers affecting clustering arise, reducing *topModNum* may help.

For the t-SNE perplexity parameter, LTR_Stream outputs the dimensionality reduction results before clustering. A too-high perplexity leads to overly dense local structures (Fig. S17D), while a too-low value results in overly dispersed clustering (Fig. S17F). The ideal perplexity setting produces clustering results as shown in Fig. S17E. We recommend an adjustment range of 50–500, with each adjustment changing the value by at least 5% of the current setting. For other parameters, refer to Table S6 and adjust as needed.

## Acknowledgements

The authors would like to thank Shujun Ou for valuable discussion.

## Data availability

Genome assemblies of three *Papaver* species and four *Gossypium* species used in this study are listed in Table S2. RNA-seq and Hi-C data of three *Papaver* species were downloaded from Sequence Read Archive (SRA) database under project accession number PRJNA720042. LTR_Stream is freely available to non-commercial users at https://github.com/xjtu-omics/LTR_Stream.

## Author contributions

K.Y. and X.Y. designed and supervised the research. T.X. wrote the codes. H.Z. tested the codes. T.X., Y.C., T.W., P.J., S.W., P.S., P.Z., S.G., Y.X., C.W., and N.D. analyzed the results. K.Y., X.Y., T.X., Y.Z., and S.B. wrote and revised the manuscript. All authors have read and approved the final version.

