## Supplementary Figures for "Deciphering complex interactions between LTR retrotransposons and three *Papaver* species using LTR_Stream"

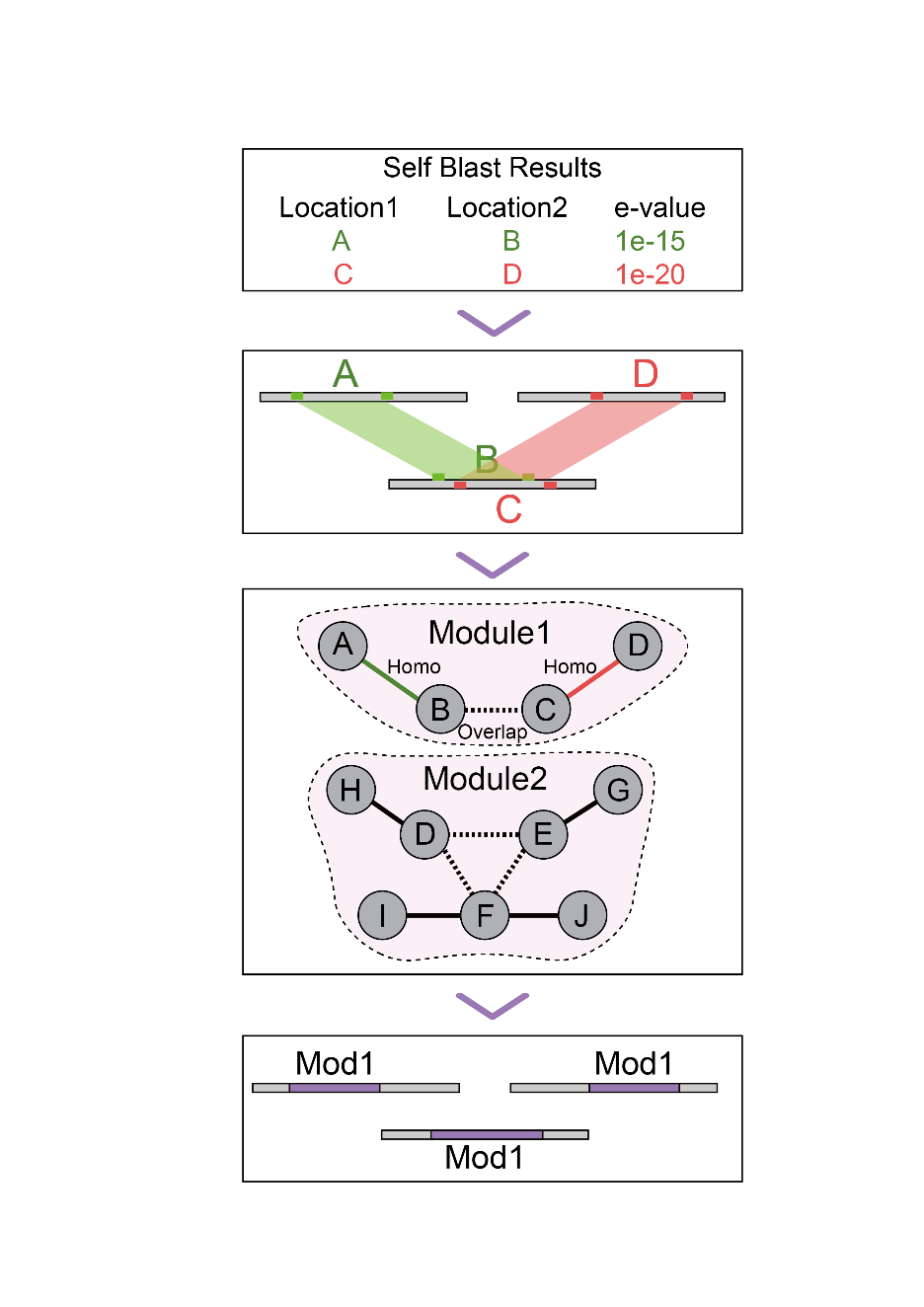

**Figure S1: Module identification in LTR_Stream.** The first panel shows samples of identified homology pairs. The second panel shows homologous pairs across different LTR-RTs (indicated by grey rectangles). The third panel shows homology graph building and connectivity identification. The solid lines indicate edges added according to homology pairs. The dashed lines indicate edges added according to overlaps. The last panel shows identified modules in LTR-RTs (marked as purple).

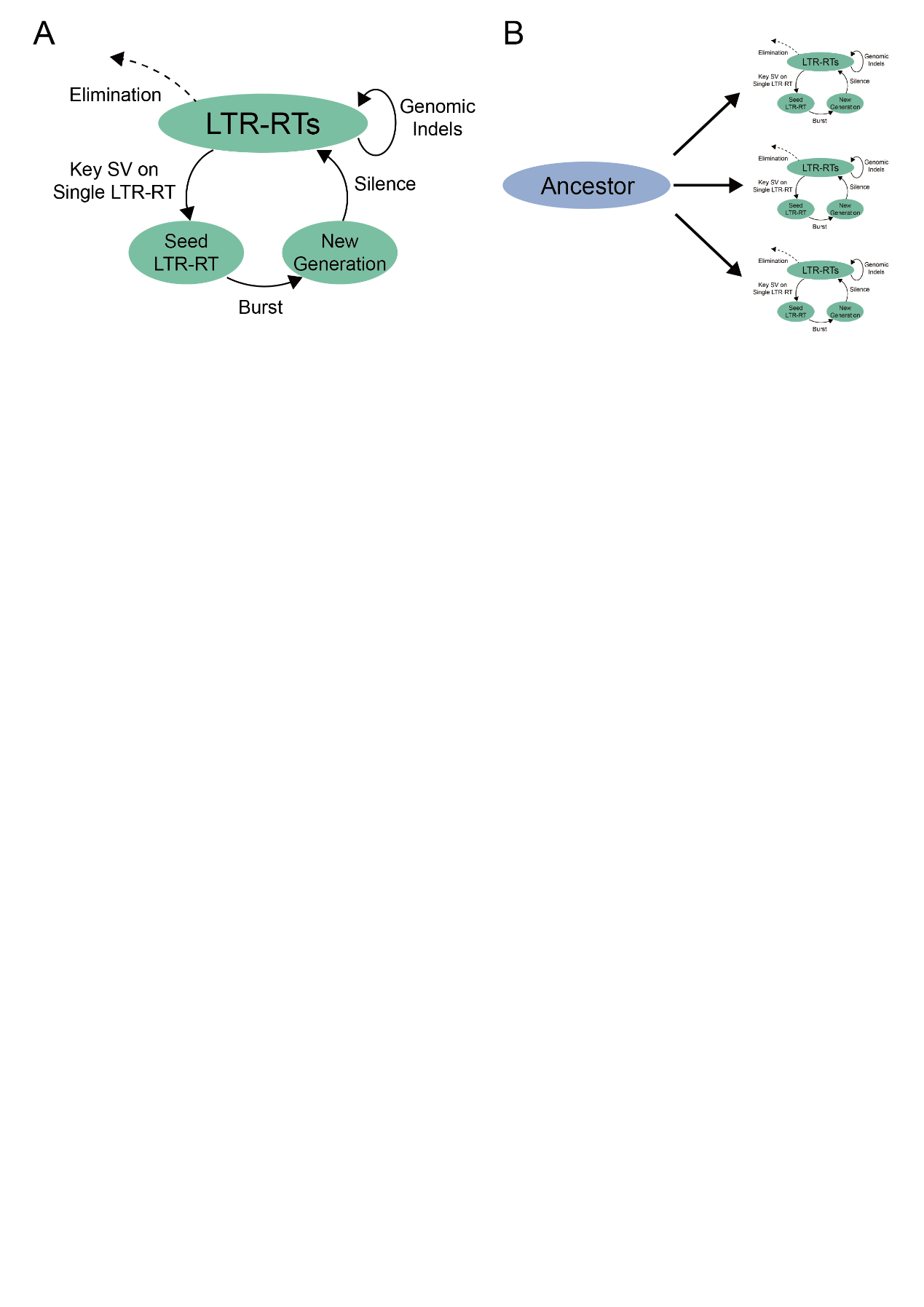

**Figure S2: Data simulation. A.** Flow chart showing simulation of one cycle of LTR-RT burst. **B.** Flow chart showing simulation of independent LTR-RT evolutionary paths.

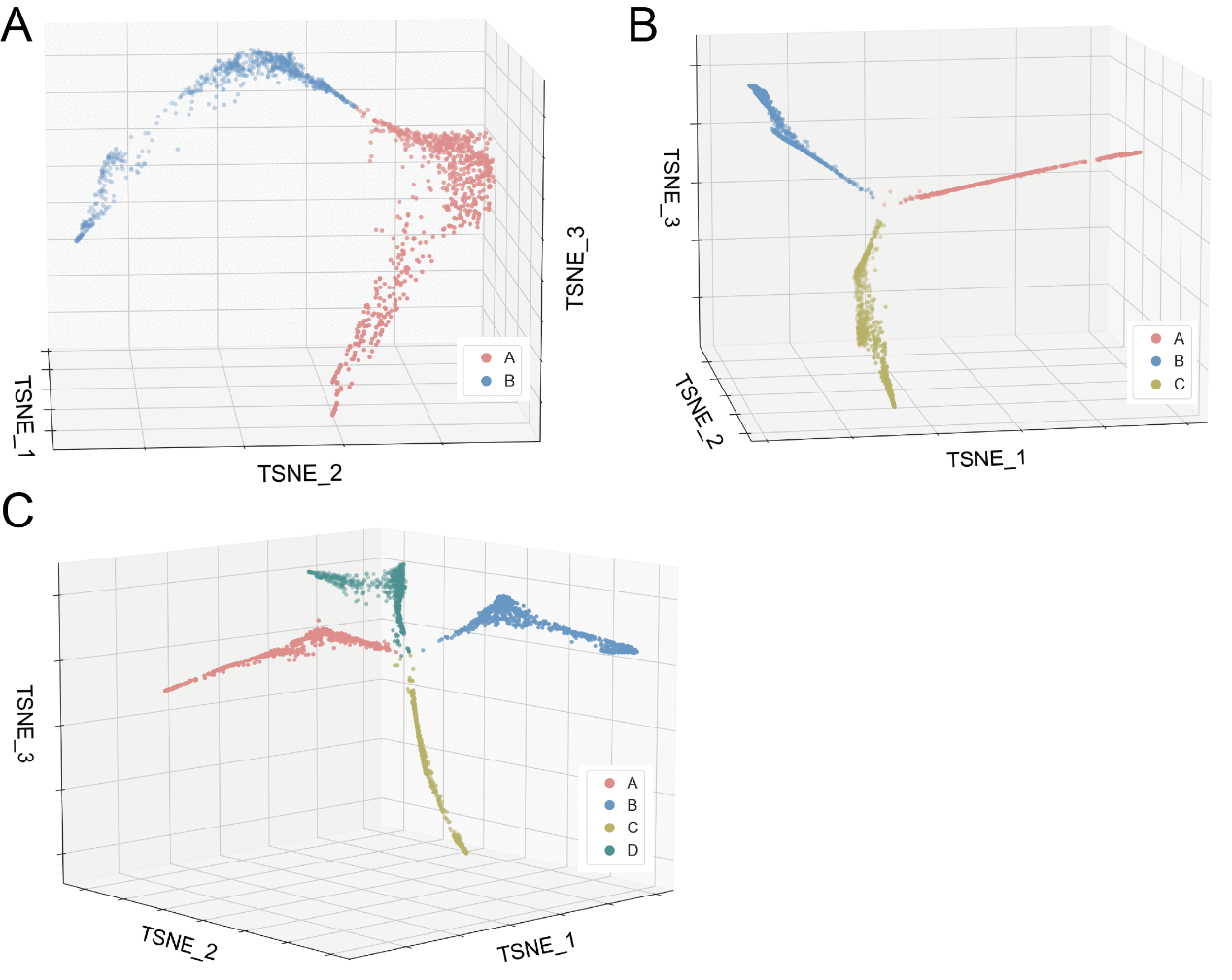

**Figure S3: LTR_Stream performance on simulated datasets based on one Ale sequence.** Panel **A, B, C** show the performance on two, three and four simulated evolutionary paths, respectively. In each panel, the left shows the reconstructed trajectories and clustering result in the 3D space. Each dot represents one module sequence that extracted from one or several LTR-RTs. Colors of dots indicate original simulated path ID.

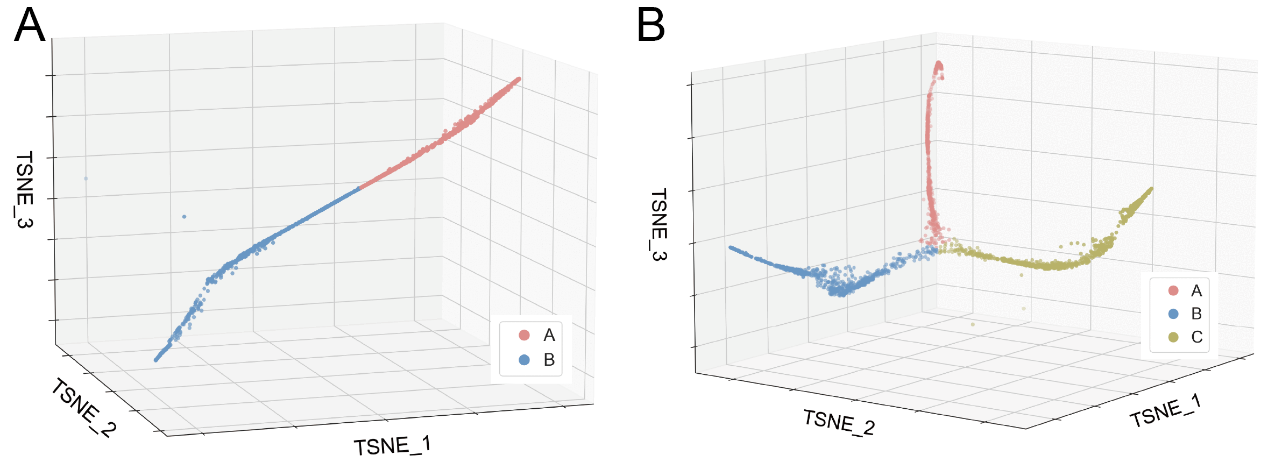

**Figure S4: LTR_Stream performance on simulated datasets based on one CRM sequence.** Panel **A, B** show the performance on two and three simulated evolutionary paths, respectively. In each panel, the left shows the reconstructed trajectories and clustering result in the 3D space. Each dot represents one module sequence that extracted from one or several LTR-RTs. Colors of dots indicate original simulated path ID.

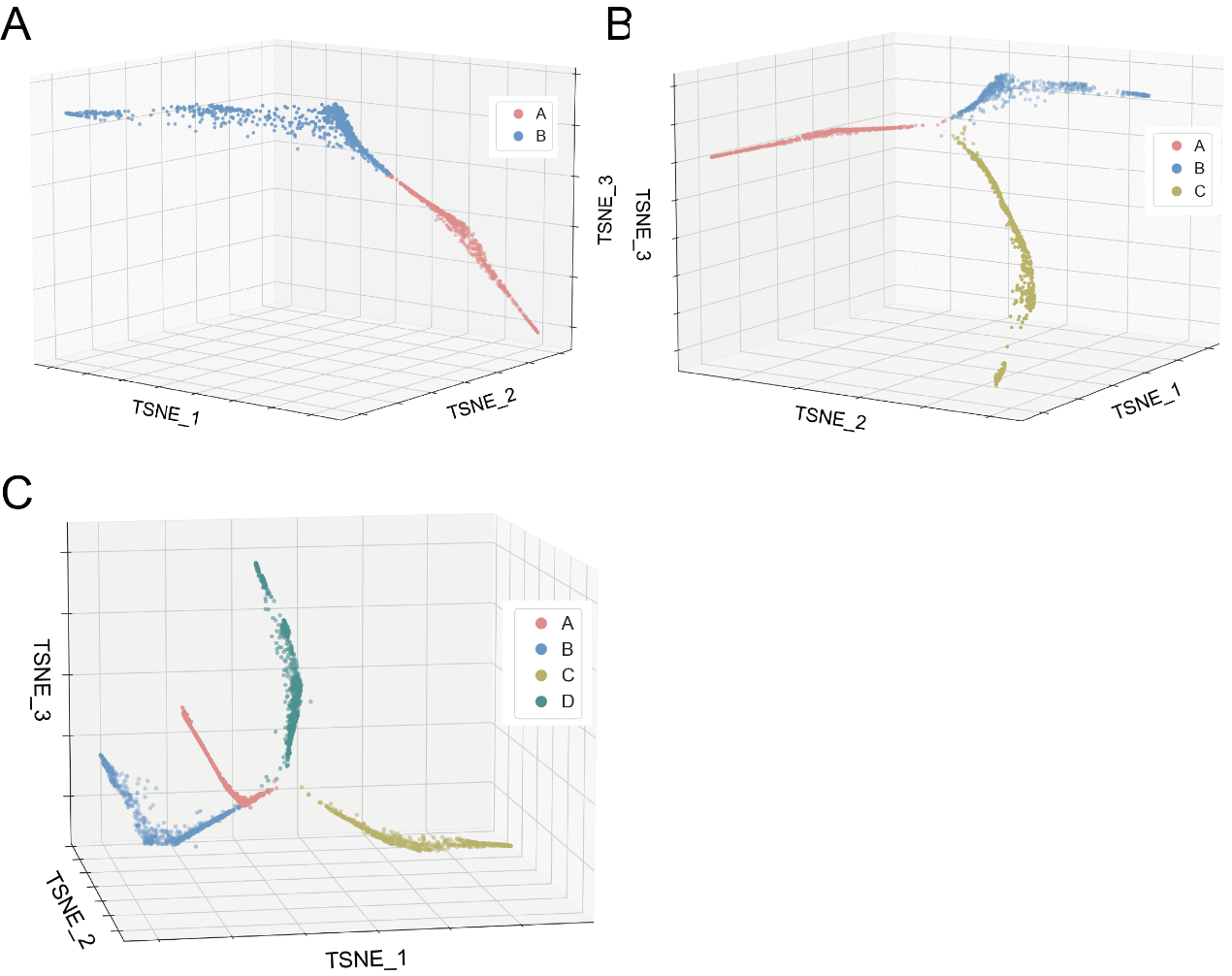

**Figure S5: LTR_Stream performance on simulated datasets based on one Tork sequence.** Panel **A, B, C** show the performance on two, three and four simulated evolutionary paths, respectively. In each panel, the left shows the reconstructed trajectories and clustering result in the 3D space. Each dot represents one module sequence that extracted from one or several LTR-RTs. Colors of dots indicate original simulated path ID.

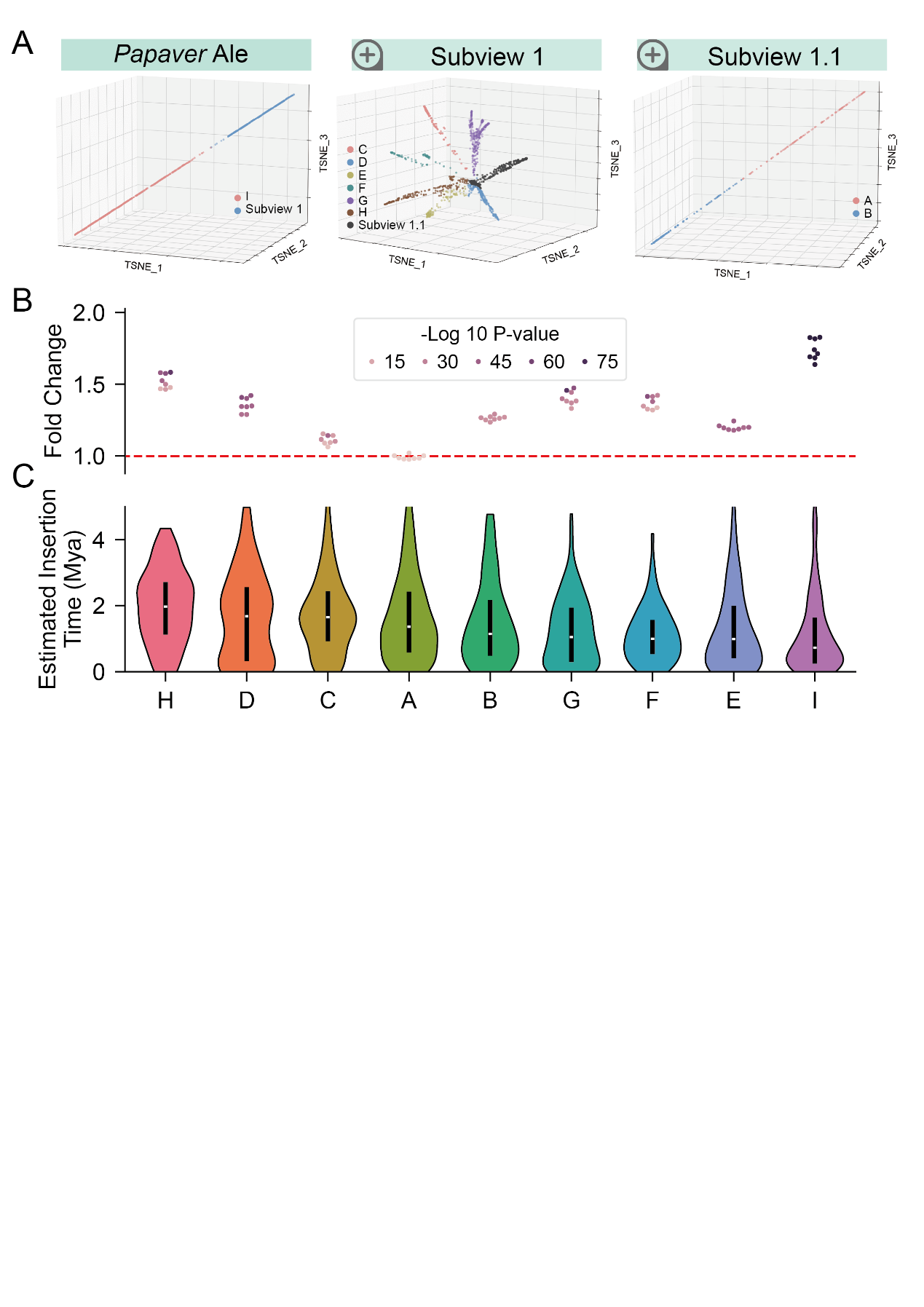

**Figure S6: Sub-lineage clustering by LTR_Stream on Ale LTR-RTs of the *Papaver* group. A.** 3D dot plots showing different sub-lineages in different subviews. Each dot represents one module sequence. Colors of dots indicate different subviews or sub-lineages. **B.** Swam plot showing fold change of inter and internal sub-lineage pairwise sequence distances. Colors of dots indicate P-value by Wilcoxon rank-sum test. The red dashed line indicates fold change value of 1.0. **C.** Violin plot showing estimated insertion time of different sub-lineages. Violins were sorted according to the median time of each sub-lineage.

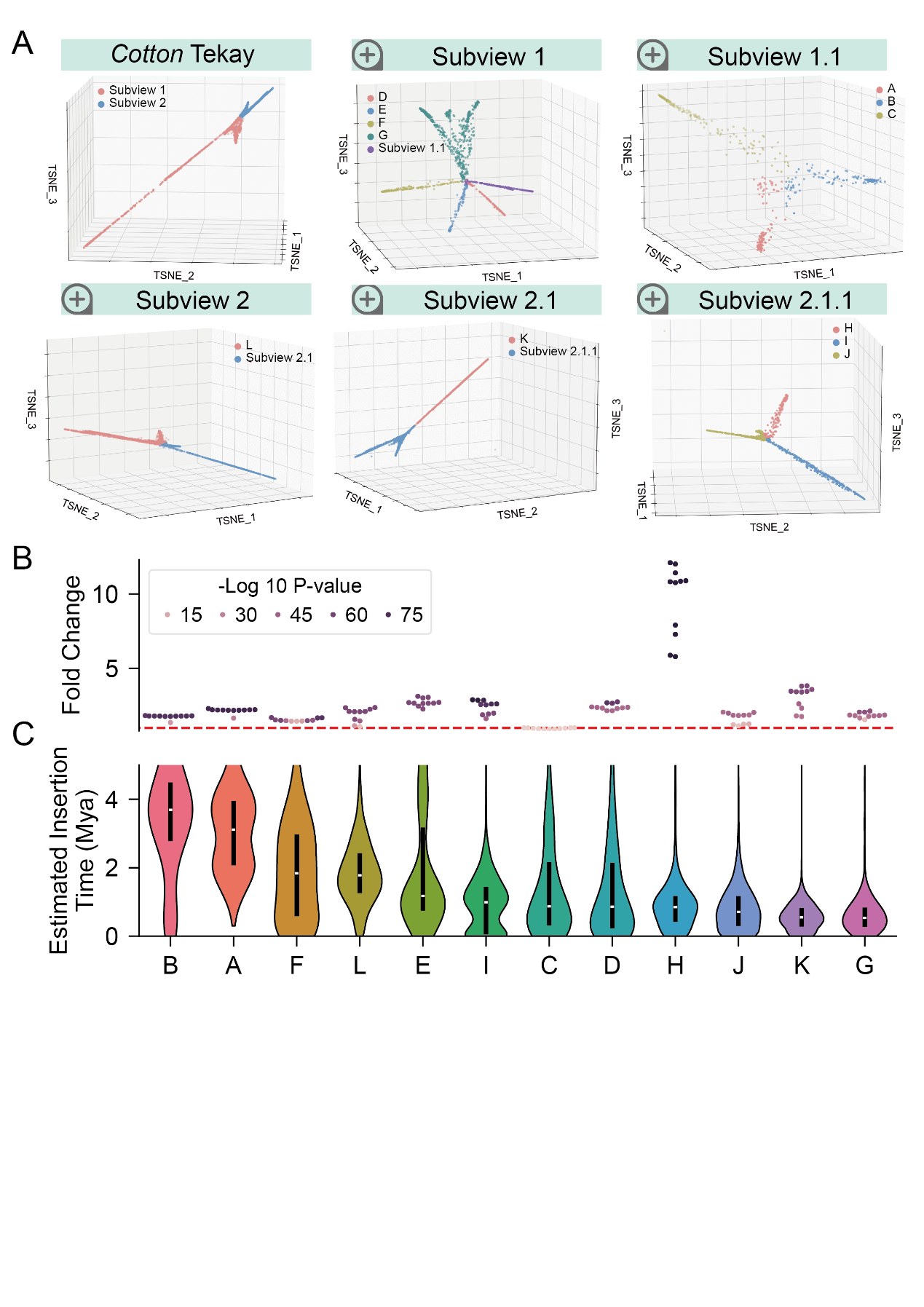

**Figure S7: Sub-lineage clustering by LTR_Stream on Tekay LTR-RTs of the *Gossypium* group. A.** 3D dot plots showing different sub-lineages in different subviews. Each dot represents one model sequence. Colors of dots indicate different subviews or sub-lineages. **B.** Swam plot showing fold change of inter and internal sub-lineage pairwise sequence distances. Colors of dots indicate P-value by Wilcoxon rank-sum test. The red dashed line indicates fold change value of 1.0. **C.** Violin plot showing estimated insertion time of different sub-lineages. Violins were sorted according to the median time of each sub-lineage.

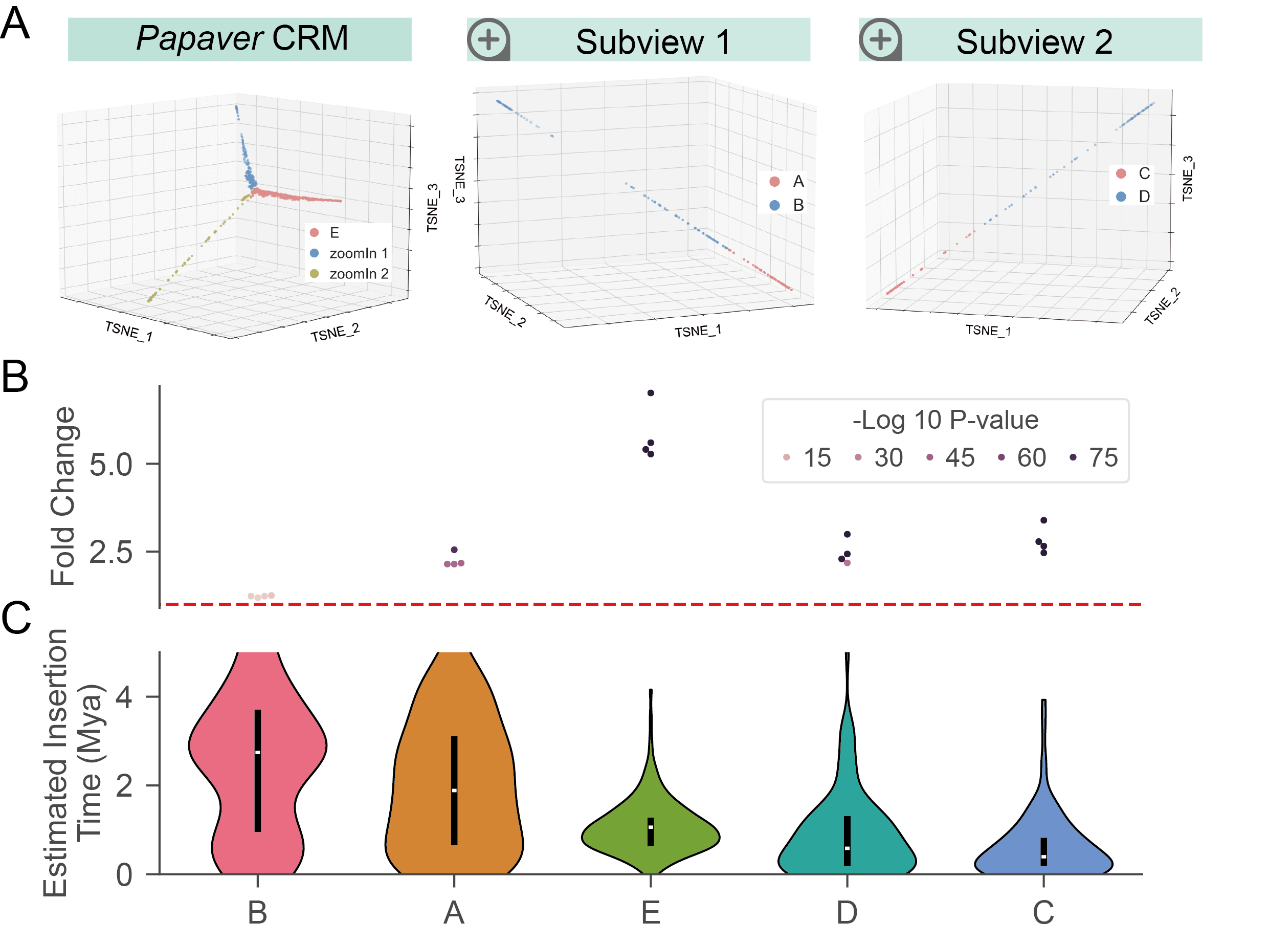

**Figure S8: Sub-lineage clustering by LTR_Stream on CRM LTR-RTs of the *Papaver* group. A.** 3D dot plots showing different sub-lineages in different subviews. Each dot represents one model sequence. Colors of dots indicate different subviews or sub-lineages. **B.** Swam plot showing fold change of inter and internal sub-lineage pairwise sequence distances. Colors of dots indicate P-value by Wilcoxon rank-sum test. The red dashed line indicates fold change value of 1.0. **C.** Violin plot showing estimated insertion time of different sub-lineages. Violins were sorted according to the median time of each sub-lineage.

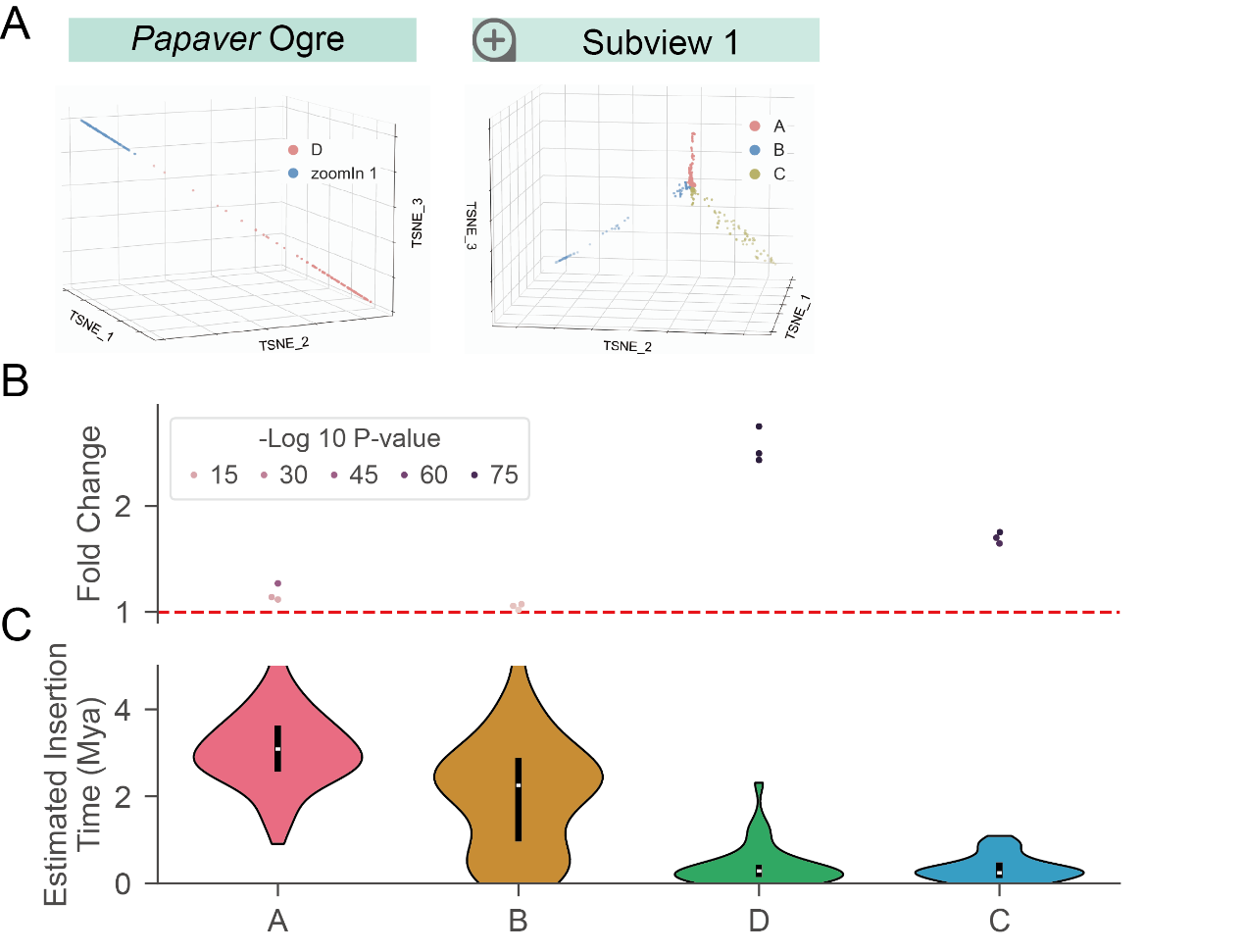

**Figure S9: Sub-lineage clustering by LTR_Stream on Ogre LTR-RTs of the *Papaver* group. A.** 3D dot plots showing different sub-lineages in different subviews. Each dot represents one model sequence. Colors of dots indicate different subviews or sub-lineages. **B.** Swam plot showing fold change of inter and internal sub-lineage pairwise sequence distances. Colors of dots indicate P-value by Wilcoxon rank-sum test. The red dashed line indicates fold change value of 1.0. **C.** Violin plot showing estimated insertion time of different sub-lineages. Violins were sorted according to the median time of each sub-lineage.

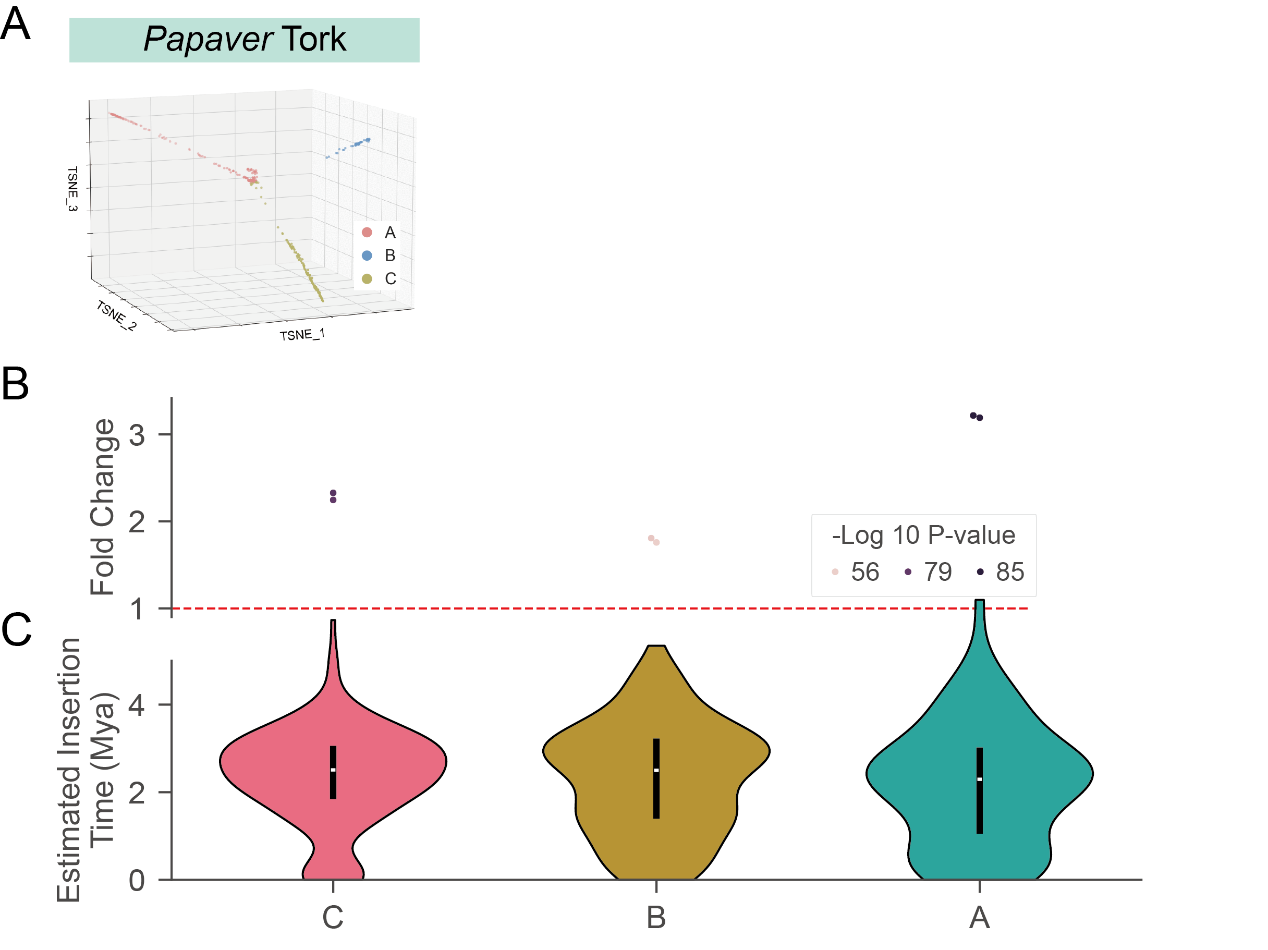

**Figure S10: Sub-lineage clustering by LTR_Stream on Tork LTR-RTs of the *Papaver* group. A.** 3D dot plots showing different sub-lineages in different subviews. Each dot represents one model sequence. Colors of dots indicate different subviews or sub-lineages. **B.** Swam plot showing fold change of inter and internal sub-lineage pairwise sequence distances. Colors of dots indicate P-value by Wilcoxon rank-sum test. The red dashed line indicates fold change value of 1.0. **C.** Violin plot showing estimated insertion time of different sub-lineages. Violins were sorted according to the median time of each sub-lineage.

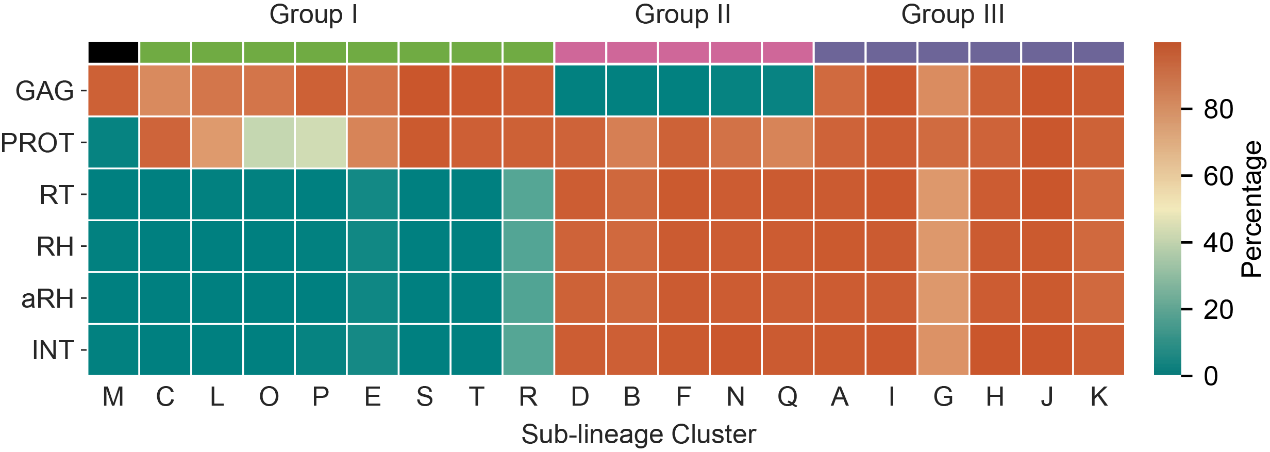

**Figure S11: Heatmap showing conserved protein domain percentage across the 20 sub-lineages of Retand. Group I:** Sub-lineages that mainly contain GAG and PROT. **Group II:** Sub-lineages that mainly contain INT, PROT, RH, RT and aRH. **Group III:** Autonomous sub-lineages that mainly contain the six conserved protein domains. Sub-lineage M, with only GAG domain, was marked as black separately. The order of protein domains (from top to bottom) follows their classical order in Retand.

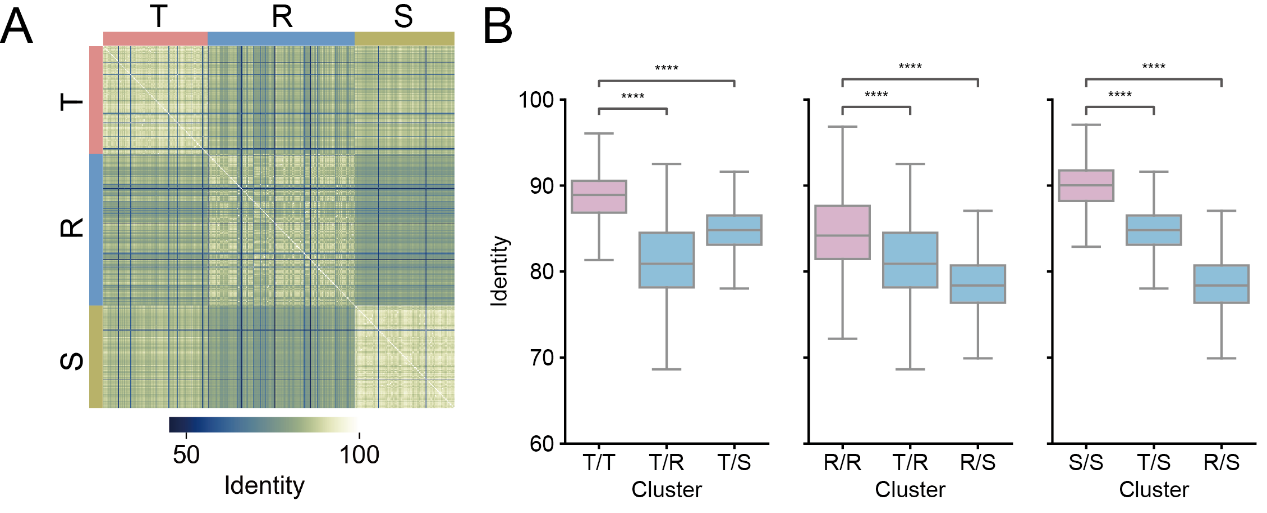

**Figure S12: Validation of sub-lineage T, R and S with nucleotide sequences. A.** Heatmap showing pairwise identities of LTR-RTs of sub-lineages T, R and S. The top and left color bars indicate their respective sub-lineage. **B.** Box plots showing pairwise inter (denoted as blue) and intra-sub-lineage (denoted as pink) identities. Differences in identity are assessed using a Wilcoxon rank-sum test, where ‘****’ indicates P-value ≤ 1e-4.

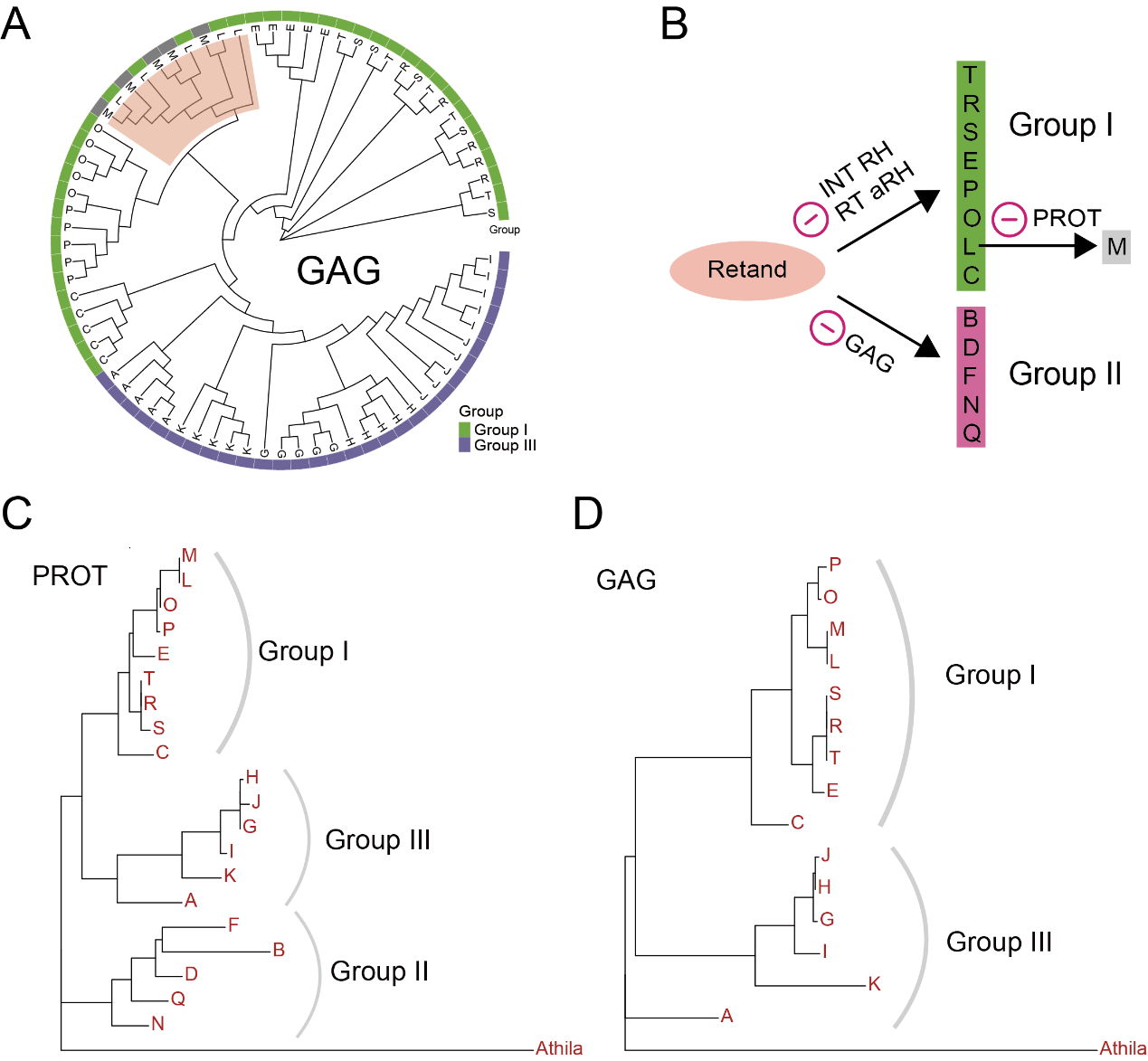

**Figure S13: A.** Rootless phylogenetic tree showing randomly sampled GAG sequences from the fifteen sub-lineages (without sub-lineages of Group II). Colors of leaf nodes indicate sub-lineages. The outermost color blocks denote their respective groups. Branch of sub-lineage L and M is denoted with pink shadow. **B.** Inferred LTR-RT non-autonomization evolutionary model. **C.** A molecular phylogenetic tree based on the consensus GAG sequences of sub-lineages. **D.** A molecular phylogenetic tree based on the consensus PROT sequences of sub-lineages. Corresponding domain sequence of Athila is used as the outgroup.

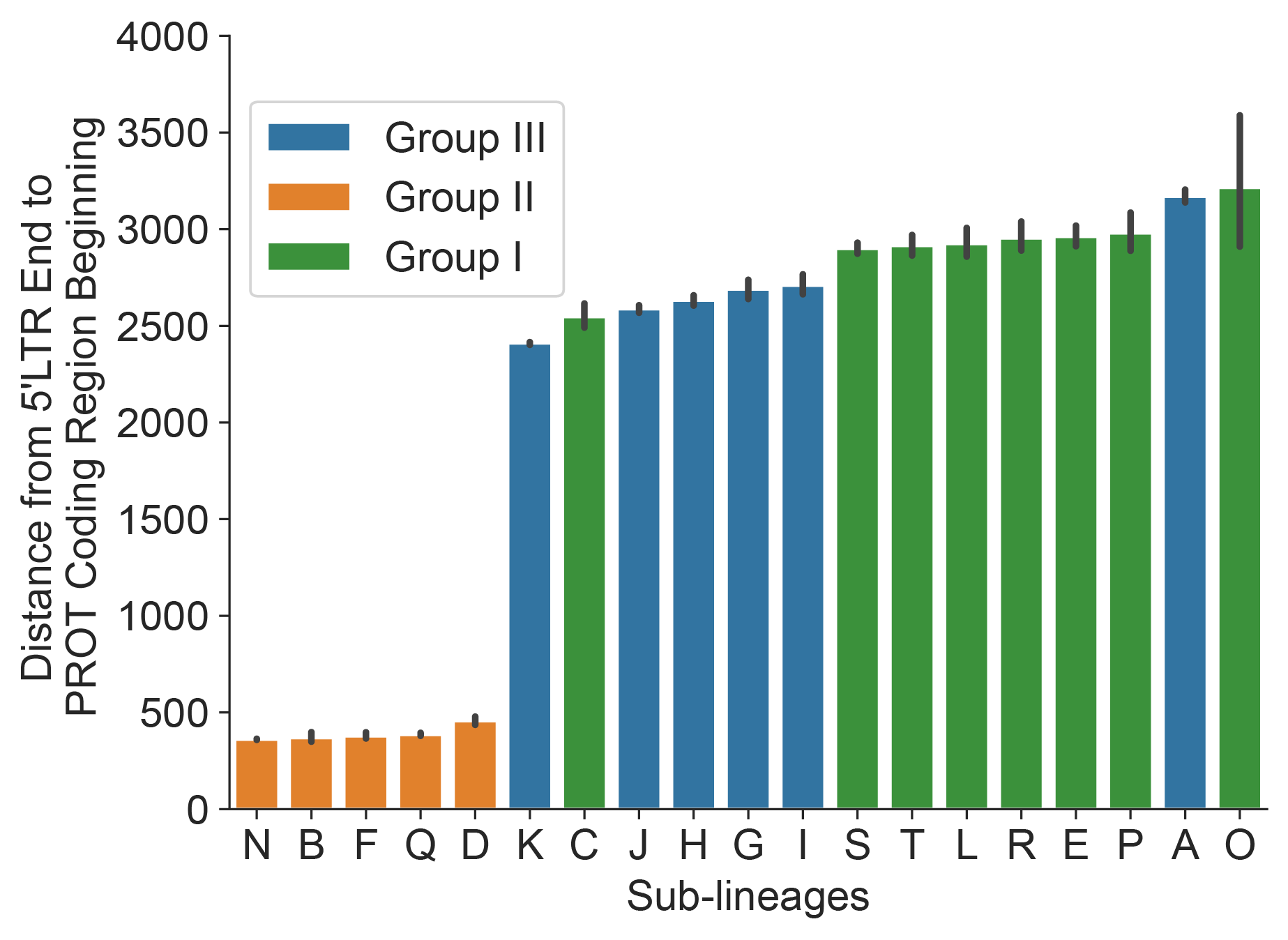

**Figure S14:** Bar plot showing distance between the 5' LTR and the protease-coding region among sub-lineages. Colors of bars indicate different Group.

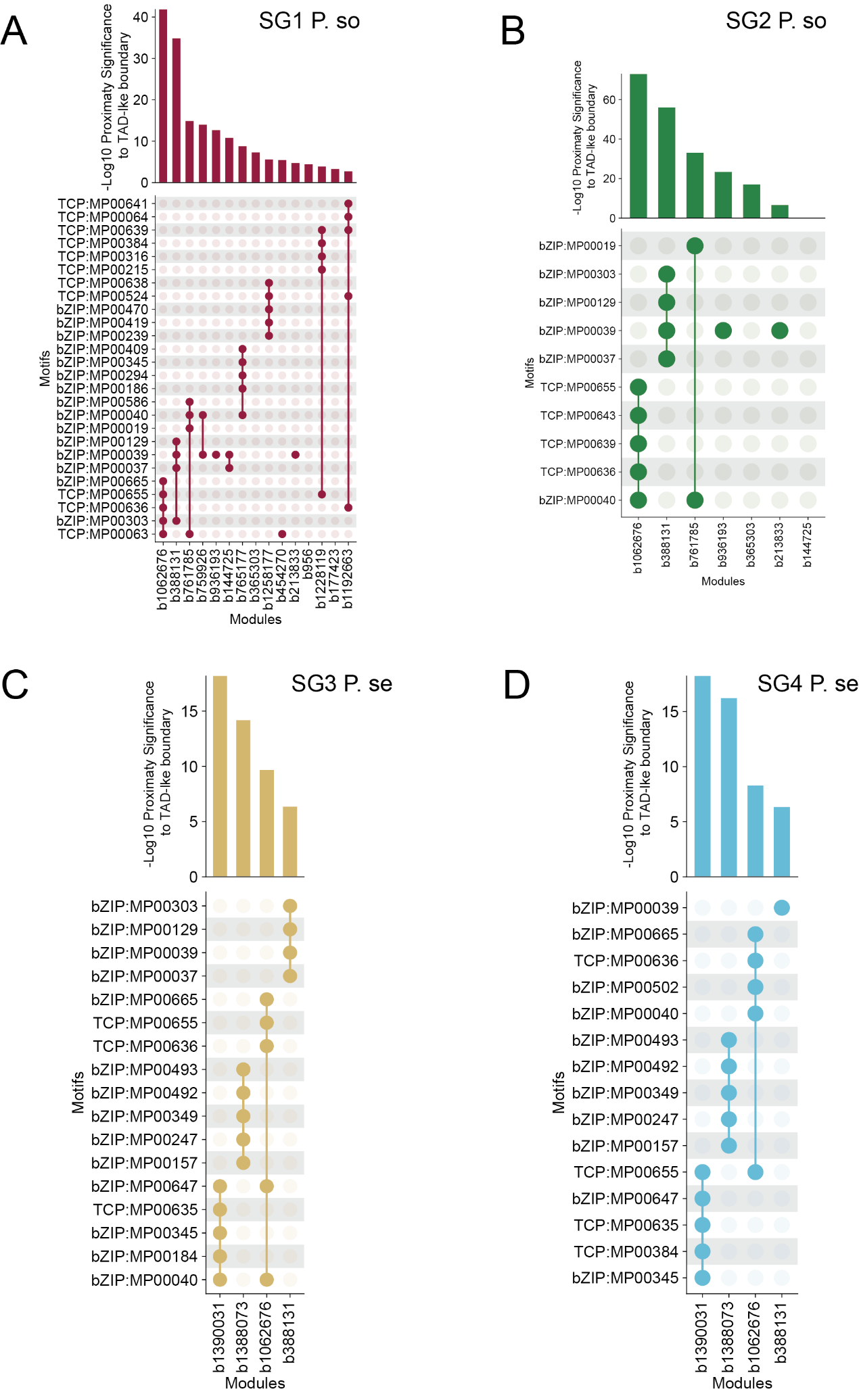

**Figure S15: Upsetplots showing enriched motifs of different module across the four subgenomes. A.** Top: Bars showing proximity significance of each module to TAD-like boundary of SG1 P. so. Bottom: Upsetplot showing enriched motifs of each module. Transcription factor type (TCP or bZIP) were annotated before motif ID. Only those motifs with P-value smaller than 1e-10 and percentage more than 70% in target sequences were displayed. **B, C and D**: data of subgenomes SG2 P. so, SG3 P. se, and SG4 P. se are presented separately in the same format.

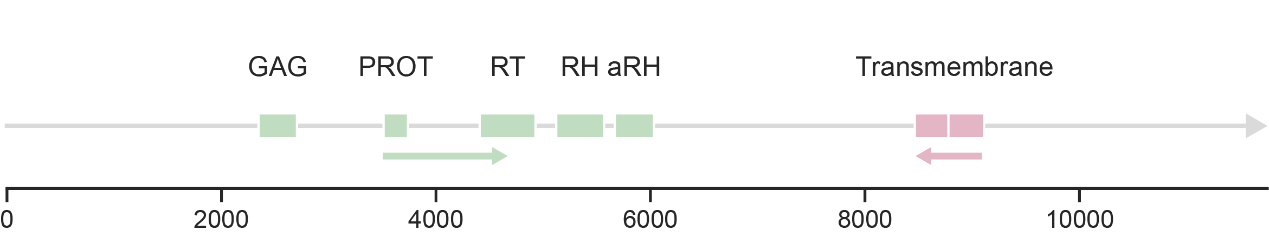
 **Figure S16: An example showing locations of ORFs of one Retand LTR-RT (located at P. se chr7:** **78938605-78950355).** Green rectangles represent ORFs of conserved proteins. Pink rectangles represent antisense ORFs that coding transmembrane proteins.

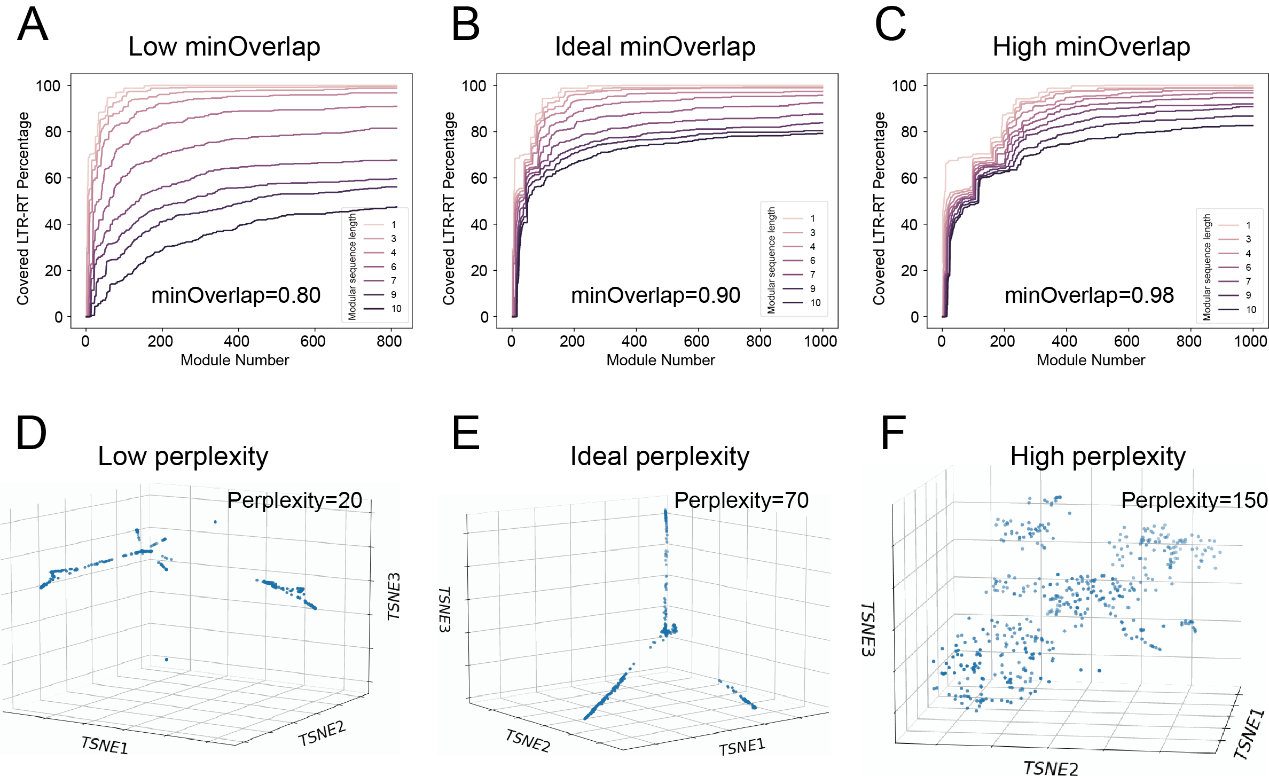

**Figure S17: Parameter adjustment guideline using the *Papaver* Tork dataset as an example. A-C.** Coverage curves for too low, ideal, and too high *minOverlap* values. Different colors represent coverage of various modular sequence lengths. **D-F.** Dot plot illustrates the dimensionality reduction results under conditions of too small, ideal, and too large *perplexity*.
