## Supplementary Table1-4 for "Deciphering complex interactions between LTR retrotransposons and three *Papaver* species using LTR_Stream"

### Supplementary Tables

#### Table S1: Simulated datasets

| **Simulated LTR-RT dataset ID** | **Ancestral LTR-lineage** | **Number of branches** | **Number of simulated LTR-RTs** |
| --- | --- | --- | --- |
| 1 | Ale | 2 | 1646 |
| 2 | Ale | 3 | 2460 |
| 3 | Ale | 4 | 3292 |
| 4 | CRM | 2 | 1631 |
| 5 | CRM | 3 | 2413 |
| 6 | CRM | 4 | 3221 |
| 7 | Tork | 2 | 1607 |
| 8 | Tork | 3 | 2382 |
| 9 | Tork | 4 | 3186 |

#### Table S2: Assemblies of closely related species used for testing LTR_Stream

| **Group** | **Species** | **Publications** | **Download Links** |
| --- | --- | --- | --- |
| *Gossypium* | *Gossypium herbaceum* | (1) | [Link 1](https://www.cottongen.org/cottongen_downloads/Gossypium_herbaceum/A1_WHU/assembly/Gherbaceum_A1-0076_WHUv3.0rc.genome.standard.fa.gz) |
|  | *Gossypium hirsutum* | (1) | [Link 2](https://www.cottongen.org/cottongen_downloads/Gossypium_hirsutum/WHU-TM1_AD1_Updated/assembly/Ghirsutum_TM-1_WHU_genome.standard.fa.gz) |
|  | *Gossypium barbadense* | (1) | [Link 3](https://www.cottongen.org/cottongen_downloads/Gossypium_barbadense/HEAU-Pima90_AD2genome/assembly/Pima90.fa.gz) |
|  | *Gossypium raimondii* | (2) | [Link 4](https://figshare.com/ndownloader/files/25304561) |
| *Papaver* | *Papaver rhoeas* | (3) | [Link 5](https://ngdc.cncb.ac.cn/gwh/Assembly/17874/show,%20GWHAZPH00000000) |
|  | *Papaver somniferum* | (3) | [Link 6](https://ngdc.cncb.ac.cn/gwh/Assembly/17875/show) |
|  | *Papaver setigerum* | (3) | [Link 7](https://ngdc.cncb.ac.cn/gwh/Assembly/17873/show) |

#### Table S3: Lineage-level classification of the LTR-RTs of the three *Papaver* species

| **LTR-RT lineage** | **No. of LTR-RTs in *P. rhoeas*** | **No. of LTR-RTs in *P. somniferum*** | **No. of LTR-RTs in *P. setigerum*** | | **Total** |
| --- | --- | --- | --- | --- | --- |
| Retand | 5915 | 2076 | 2208 | 10199 | |
| Ale | 798 | 1324 | 1999 | 4121 | |
| Athila | 124 | 2133 | 1657 | 3914 | |
| Unknown | 790 | 924 | 866 | 2580 | |
| Ivana | 337 | 786 | 949 | 2072 | |
| CRM | 506 | 478 | 610 | 1594 | |
| Reina | 497 | 469 | 595 | 1561 | |
| TAR | 63 | 411 | 688 | 1162 | |
| Bianca | 329 | 214 | 363 | 906 | |
| Ogre | 353 | 167 | 192 | 712 | |
| Tork | 203 | 172 | 252 | 627 | |
| Ikeros | 52 | 86 | 107 | 245 | |
| mixture | 133 | 13 | 23 | 169 | |
| SIRE | 30 | 30 | 33 | 93 | |
| Alesia | 19 | 11 | 27 | 57 | |
| Tekay | 34 | 6 | 7 | 47 | |
| Galadriel | 21 | 7 | 8 | 36 | |
| Angela | 2 | 0 | 2 | 4 | |
| Gymco-III | 1 | 0 | 0 | 1 | |
| Osser | 1 | 0 | 0 | 1 | |
| No Result | 2417 | 3537 | 2966 | 8920 | |
| Total | 12625 | 12844 | 13552 | 39021 | |

#### Table S4: Lineage-level classification of the LTR-RTs of the four *Gossypium* species

| **LTR-RT lineage** | **No. of LTR-RTs in *G. barbadense*** | **No. of LTR-RTs in *G. herbaceum*** | **No. of LTR-RTs in *G. hirsutum*** | **No. of LTR-RTs in *G. raimondii*** | **Total** |
| --- | --- | --- | --- | --- | --- |
| Tekay | 3208 | 4967 | 4407 | 48 | 12630 |
| Tork | 1854 | 645 | 2098 | 243 | 4840 |
| CRM | 1097 | 109 | 1370 | 125 | 2701 |
| Ivana | 864 | 163 | 866 | 212 | 2105 |
| Ale | 654 | 308 | 636 | 281 | 1879 |
| unknown | 270 | 220 | 524 | 4 | 1018 |
| Athila | 313 | 285 | 295 | 54 | 947 |
| Galadriel | 286 | 78 | 305 | 127 | 796 |
| Ogre | 210 | 106 | 236 | 12 | 564 |
| Reina | 88 | 45 | 85 | 43 | 261 |
| TAR | 114 | 14 | 104 | 27 | 259 |
| Bianca | 87 | 58 | 87 | 14 | 246 |
| Ikeros | 36 | 32 | 36 | 3 | 107 |
| Angela | 38 | 8 | 33 | 11 | 90 |
| Alesia | 12 | 5 | 9 | 5 | 31 |
| mixture | 5 | 1 | 2 | 0 | 8 |
| SIRE | 0 | 0 | 1 | 1 | 2 |
| Selgy | 0 | 1 | 0 | 0 | 1 |
| No Result | 283 | 516 | 331 | 32 | 1162 |
| Total | 9419 | 7561 | 11425 | 1242 | 29647 |
